# High-throughput two-photon volumetric brain imaging in freely moving mice

**DOI:** 10.1101/2024.10.13.618106

**Authors:** Long Qian, Yaling Liu, Yalan Chen, Jianglai Wu

## Abstract

Imaging neural activities across large volumes at subcellular resolution without impeding animal behaviors remains difficult. Here, we develop a high-throughput miniature Bessel-beam two-photon microscope (miniBB2p) capable of imaging calcium dynamics from neurons and dendrites over a volume of 420 × 420 × 80 μm³ during free behavior. We share a full description of miniBB2p and demonstrate its ability to perform large-scale recordings of the calcium activities of more than 1000 neurons at a time in anterior cingulate cortex and secondary motor cortex of freely moving mice in various behavioral paradigms. Our results indicate that miniBB2p opens new avenues in the design of miniature multiphoton microscopes, facilitating precise monitoring of neural activities across large populations within a three-dimensional volume in freely moving animals.

---

Head-mounted miniature multiphoton microscopy offers a promising avenue for investigating neural activity during free and naturalistic behavior in mice and rats, thanks to its inherent optical sectioning and deep tissue penetration capabilities [1]. Recent advancements in optical fibers, miniature detectors, laser scanners, focus-tunable lenses, and objectives have significantly improved the performance of miniature multiphoton microscopes [2–10], leading to a substantial increase in the number of neurons that can be imaged in freely moving animals in practical neurobiological investigations [10–13]. However, several limitations are presented in existing miniature multiphoton microscopes. First, rapid volumetric imaging is challenging due to the sequential acquisition of each voxel, especially when using high numerical aperture (NA) objectives that offer superior lateral and axial resolution. While advanced platforms with fast focus-tunable lenses can scan the focus axially in milliseconds [5,10], a restricted number of thin optical sections within a volume can be imaged at a time, albeit at the expense of imaging speed and resolution. Second, the shallow depth of field of the microscope makes imaging susceptible to brain tissue movement, particularly axial movement. This issue is a common concern in imaging head-fixed animals [14] and would be exacerbated when imaging freely moving animals as brain movement is less constrained [8]. Finally, while customized optics such as miniature objectives have been utilized to achieve higher resolution and larger field of view (FOV), achieving diffraction-limited resolution across the FOV remains difficult [6, 8, 10]. This challenge arises from the need to fit all optics into a compact space of a few cubic centimeters, which poses extra hurdles for lens design and assembly.

Bessel beam can address these limitations. In recent years, tabletop two-photon microscopes (2PM) utilizing Bessel beams have shown several distinct features over the widely used Gaussian beams [15–20]. Gaussian-beam 2PM records a stack of 2D optical sections across the depth to achieve 3D imaging, it favors resolving the fine structures in the axial dimension at a compromised imaging speed. In contrast, Bessel-beam 2PM captures projected views of volume structures by scanning an axially elongated focus in 2D. This process transfers the 2D scan rate into a volumetric imaging rate, increases the information throughput, and reduces the data size. It is an effective tool for volumetric calcium imaging in vivo, as the position of neurons and their fine structures remain stable during imaging [17]. In addition, if needed, the axial coordinates of structures in the 2D projections can be retrieved with a full 3D reference image from a Gaussian-beam 2PM. The extended depth of field also makes imaging resistant to axial movements of brain tissue [17], which is preferred in monitoring neural circuits in freely moving animals. Moreover, in comparison to Gaussian-beam 2PM, Bessel-beam 2PM achieves better lateral resolution at same excitation NA [17–18] and is mainly sensitive to astigmatism among all primary optical aberrations, indicating the potential of designing simple lenses for miniature microscopes [20–21]. In this work, taking advantage of these unique features of Bessel beam in two-photon imaging, we developed a head-mounted miniature Bessel-beam two-photon microscope (miniBB2p) and demonstrated high-throughput volumetric calcium imaging in secondary motor cortex (M2) and in anterior cingulate cortex (ACC) in freely moving mice.

## RESULTS

### Bessel beam simplifies the optical design in miniBB2p

Among all primary optical aberrations, including spherical aberration, coma, astigmatism, field curvature and distortion, Bessel beam is mainly sensitive to astigmatism [20–21]. Leveraging this characteristic, we utilized a systematic approach in miniBB2p to eliminate astigmatism while minimizing other primary aberrations. Briefly, we adopted a symmetric design for the scan and tube lenses, each comprising two identical plano-convex lenses (**Fig. 1a**). This symmetry minimizes coma, distortion, and lateral color aberration. Uncorrected astigmatism was fully compensated by the objective formed by two plano-convex lenses (**Fig. 1b**). In parallel, other aberrations in the objective were well-contained under the low excitation NA of 0.3, leading to near-diffraction-limited performance over a FOV of Ø 600 μm (**Fig. 1c**). Throughout the entire FOV, we found negligible blurring of the Bessel focus from the residual spherical aberration and coma, in contrast to the blurring observed with a Gaussian focus (**Supplementary Fig. 1**). We noted that the residual field curvature and distortion simply shift the focal position without blurring the focus (**Fig. 1d and Supplementary Fig.1**). The systematic design approach focused on astigmatism correction in miniBB2p significantly reduced the number of lenses in the microscope compared to conventional designs that seek to correct all major aberrations for individual lenses. This reduction not only lowered manufacturing costs but also simplified assembly and alignment, while still maintaining high image quality (**Supplementary Table 1**).

**Fig. 1.**
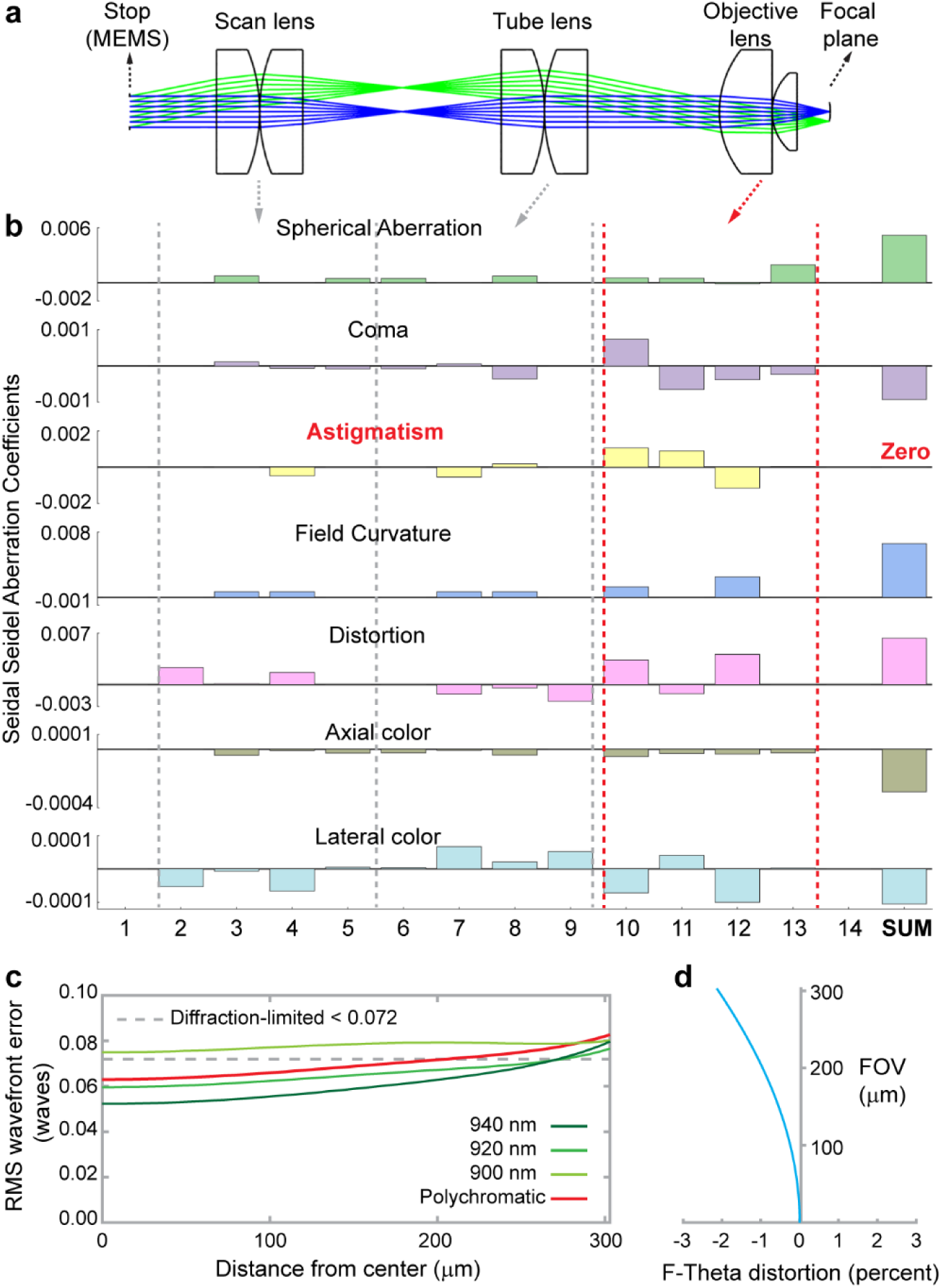
Optical design in miniBB2p. **a,** Unfolded excitation beam path from the MEMS mirror (stop) to focal plane. Blue and green rays represent the beam focusing at the center and edge of the FOV; the focal plane is curved with a radius of 2 mm, resulting in a maximum focal shift of ∼10 μm across the entire FOV. **b**, Seidel aberration coefficients across lenses. **c**, Root-mean-square wavefront error throughput a FOV of Ø 600 μm. d, Distortion of the microscope; maximum distortion is ∼ -2.1%. The stop size is 1 mm and NA is 0.3 in the simulation (See optical design files in **Supplementary Data)**.

### System design and performance testing of miniBB2p

The headpiece of miniBB2p weighed a total of 2.6 grams and was about 2.4 × 0.9 × 2.0 cm^3^ in size, making it well suited for brain imaging in freely moving mice (**Figs. 2a-d, Supplementary Fig. 2 and Methods**). We used a 1-meter large-mode-area photonic crystal fiber (LMA-12, NKT Photonics) to transmit the 920-nm femtosecond laser pulses for two-photon excitation. This endlessly single-mode photonic crystal fiber supports single-mode delivery over a broad spectral range from 700 to 1700 nm [22]. The pulse in the fiber was broadened by material dispersion and nonlinear effects (e.g., self-phase modulation) [23]. Benefitting from the large mode area (∼ Ø 10 µm), nonlinear effects in the fiber were substantially reduced. The pulse width after dispersion precompensation at the fiber output end was less than 200 fs at laser powers below 40 mW and less than 300 fs at laser powers below 100 mW (**Supplementary Fig. 2b and Methods**). To generate the Bessel beam, we engineered a custom micro-axicon with an apex angle of 120° at the fiber tip **(Fig. 2b, Supplementary Fig. 2c and Methods)**. A microelectromechanical systems (MEMS) mirror was used as the scanner, enabling a maximum square FOV of 420 × 420 μm^2^ and a frame size of 512 × 512 pixels at 9 frames per second.

**Fig. 2.**
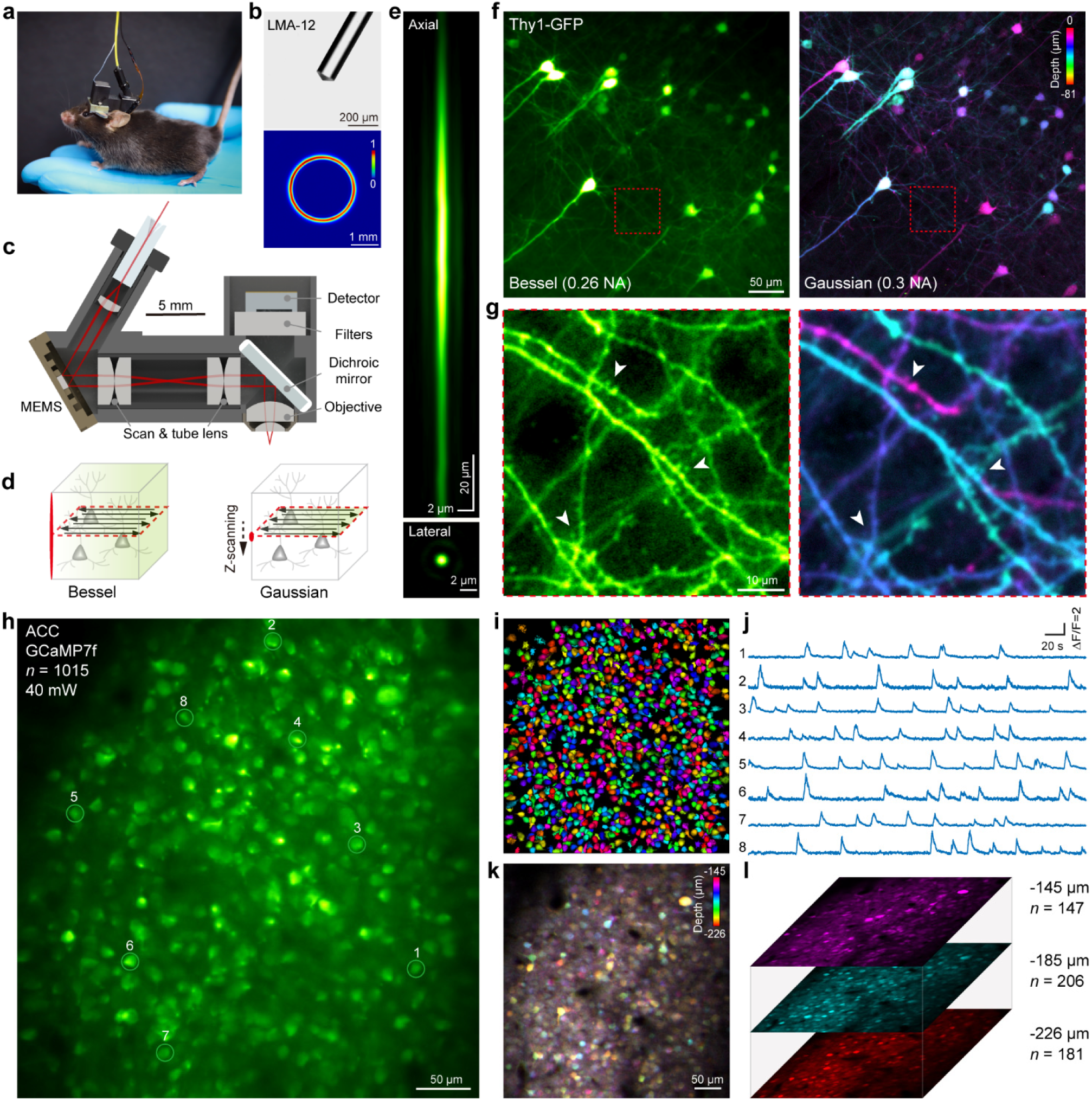
System design and test of miniBB2p. **a**, Photograph of a mouse wearing the microscope headpiece. **b**, Brightfield micrograph of the custom micro-axicon at the LMA-12 fiber tip (top) and ring pattern imaged at ∼5-mm distance from the fiber tip (bottom). **c**, Schematic of miniBB2p showing key elements and excitation beam path (red lines). **d**, Imaging a volume requires a single 2D scan of a Bessel focus but a stack of 2D scans of a Gaussian focus. **e**, Lateral and axial point spread functions measured with 500-nm fluorescent beads. **f**, Image of a ∼80-μm thick volume of Thy1-GFP-M brain slice by miniBB2p (left) and an image stack at 3-μm step collected by a tabletop Gaussian beam 2PM from the same volume, with structures colored by depth (right). **g**, zoom-in imaging of the boxed areas in **f**; white arrowheads denote dendritic spines that were more clearly visualized by Bessel focus. **h**, Imaging the spontaneous activity of neurons in anterior cingulate cortex (ACC) expressing GCaMP7f in a free-moving mouse with miniBB2p. **i**, Spatial distribution of 1015 neurons extracted by *Suite2P* in **h**, randomly colored for visualization. **j**, Example calcium signals from neurons circled in **h**. **k-l**, Imaging the same volume in awake head-fixed mouse by the tabletop 2PM: Gaussian imaging stack at 3-μm step with structures colored by depth (**k**); neural activity imaging in the top, middle, and bottom layers of the volume, the imaging depth and number of neurons identified by *Suite2P* are listed at right (**l**). Data in **h** are from a single representative experiment (n=5 animals, one experiment per animal).

The miniBB2p featured a working distance of 1 mm in water, an excitation NA of 0.26, and a collection NA of 0.95. Lateral and axial resolution measured with 500-nm fluorescent beads were 0.97–0.99 μm and 80–90 μm across the FOV (**Fig. 2e and Supplementary Fig. 3).** The lateral resolution obtained is equivalent to the diffraction-limited resolution using a Gaussian beam at a higher NA of 0.35. Notably, fluorescence collection efficiency is 2–4 times greater than that in the state-of-the-art platforms with a comparable FOV and lateral resolution [8–10] (**Supplementary Figs. 4a-d and Supplementary Table 1**). In addition, fluorescence was directly detected after the objective and filters using an onboard silicon photomultiplier, which offered higher detection efficiency and less tethered burden than fluorescence collection using thick fibers as a relay unit [6].

### Imaging comparison between miniBB2p and tabletop Gaussian-beam 2PM

We first imaged Thy1-GFP-M mouse brain slices to demonstrate the throughput and resolution of miniBB2p (**Figs. 2f-g**). Using a commercial tabletop Gaussian-beam 2PM (0.3 NA, axial resolution 22.9 μm), a stack of at least 4 images is required to cover all structures within an 80-μm thick volume. In comparison, miniBB2p captured the same information in one image, with a fourfold increase in depth of field. Moreover, the higher resolution in Bessel focus allowed easier visualization of dendritic spines (**Fig. 2g**). These observations were also found in head-fixed mouse brain **(Supplementary Fig. 5)**. We then employed miniBB2p to image spontaneous neural activities in freely moving mice. We resolved calcium dynamics of 1015 neurons at a time in ACC **(Figs. 2h-j and Supplementary Video 1)**. This result was validated through structural imaging at same region and neural activity imaging in the top, middle, and bottom layers within same volume using the tabletop 2PM on head-fixed mice (**Figs. 2k-l**). The latter approach identified 534 neurons across the three layers, approximately half the number of neurons resolved with miniBB2p. Beyond that, miniBB2p was able to image calcium transients in dendritic trees and spines along multiple dendritic branches in an axially extended volume (**Supplementary Figs. 6-7 and Supplementary Videos 2-4**). These measurements underscored the benefits of high-resolution volumetric imaging enabled by miniBB2p.

### MiniBB2p enables longitudinal recordings with minimal constraints on animal behavior

To assess the impact of carrying miniBB2p during imaging on animal behavior and mobility, we compared the movements between the naive mice carrying nothing and the mice carrying the dummy microscope with the same weight, fiber and cables as miniBB2p (**Fig. 3 and Methods**). While the mice freely exploring the behavioral arena, no significant difference were found in the total distance traveled (**Figs. 3b and d**), mean and median running speed (**Figs. 3c and e**), and the number of entries into the central coverage (**Figs. 3a and f**) between the naive mice and mice with dummy. In addition, we did not observe any notable variations of the running distance in the central and corner area of the arena following installation of the dummy microscope (**Fig. 3d**). These observations suggested that carrying miniBB2p along with the fiber and cables did not significantly impede the animal behavior. In addition, we evaluated the feasibility of conducting longitudinal studies using miniBB2p by imaging the same FOV and examining the number of identical cells recorded over multiple days. Our results revealed that the same neuronal populations could be monitored for weeks, with minimal photobleaching and photodamage (**Supplementary Figs. 8-10**), highlighting miniBB2p’s suitability for chronic imaging applications.

**Fig. 3.**
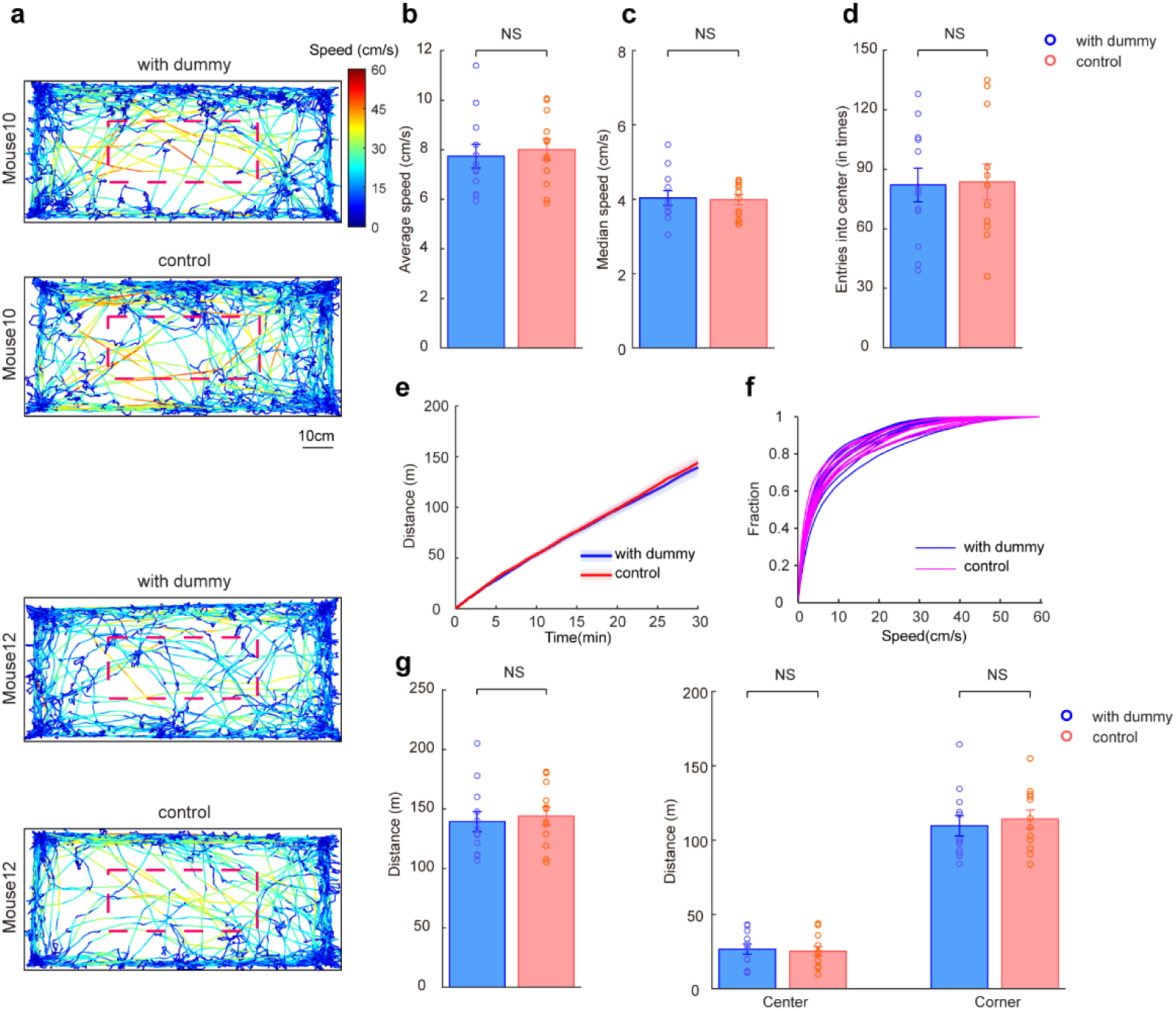
Minimal impact of miniBB2p on animal movement. **a**, Representative trajectories of mice running in a 97.6 cm × 45 cm behavioral arena in a 30-min trail; with dummy, mice carrying the 2.6 g dummy microscope, fiber, and cables; control, mice carrying nothing; running speed is color coded; red dashed box indicates the central area. **b-c**, Average speed (**b**) and median speed (**c**) in each condition (30 min each); p = 0.4156 and 0.7429, paired t test. **d**, Number of entries into the central area. p = 0.8543, paired t test. **e**, Mean accumulated distance over the 30-min test, shaded areas show s.e.m. **f**, Cumulative speed distribution over the 30-min test among all trails. **g**, Total distance traveled (left), distance traveled in the central and corner area (right) in each condition (30 min each); p = 0.4141, 0.6408 and 0.3394, paired t test. NS, no significant; data in **b-d** and **g** are presented as mean ± s.e.m. Data represent measurements collected from 12 individual animals.

### Using miniBB2p for high-throughput imaging of the neural activities across different behavioral paradigms

We used miniBB2p to record high-throughput proof-of-principle data from neuronal populations in M2 and ACC of freely moving mice in a variety of behavioral paradigms. First, we investigated the response of pyramidal neurons in M2 to locomotion over a 30-minute recording period in freely moving mice. (**Figs. 4a-c and Supplementary Video 5**). Running increased the mean ΔF/F in most neurons (**Fig. 4d**), with a running modulation index (MI) showing increased activity during running (**Fig. 4e**; n=878 neurons, running MI 0.156 ± 0.006, mean ± s.e.m.). In addition, we inspected the average fluorescence changes aligned with running onsets and found that 38.2% of the neurons recorded significantly increased their activities upon running onset (**Fig. 4f and Methods**), resembling the study from recordings in primary somatosensory cortex (S1) [24].

**Fig. 4.**
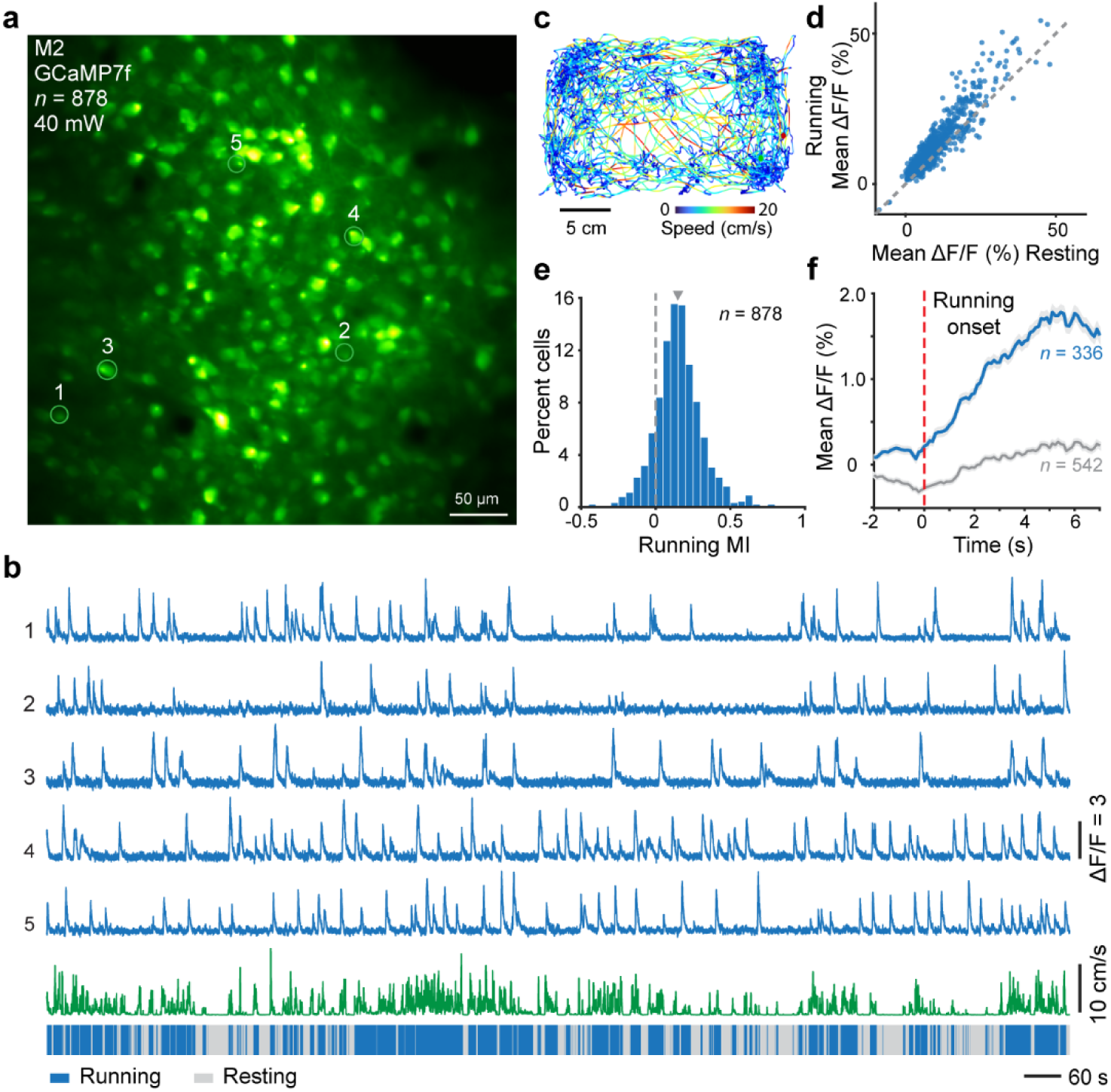
Imaging the response of pyramidal neurons in M2 to locomotion. **a**, Image of an example neuronal population in M2. **b**, Representative calcium traces along with moving speed from neurons circled in **a**; running and resting states are color coded with blue and gray. **c**, Movement trajectory and speed of the mouse over a 30-min recording period. **d**, Scatter plot of the mean ΔF/F values in resting and running states for the 878 neurons recorded; dashed line indicates a unity line. **e**, Distribution of running modulation index for the neurons; triangle indicates median. **f**, Population average of ΔF/F traces for neurons with and without increasing calcium activity in response to the running onset (16 events), shaded areas indicate s.e.m. Data represent one of three experiments in one animal.

Next, we examined the neural activity in ACC under different pain stimulation paradigms in freely moving mice. We imaged the calcium activity of 673 ± 104 (5 mice, mean ± s.e.m.) pyramidal neurons both before and after administrating a peripheral stimulus, either a noxious 27 G pin prick or control, non-noxious 0.4g von-Frey filament, to the paw contralateral to the recording site of the brain (**Figs. 5a-b, Supplementary Video 6 and Methods**). After signal processing and identification of active neurons (**Fig. 5c**), we observed that ACC neurons exhibited only low-level calcium activity in response to non-noxious stimuli. In contrast, these neurons showed an increase in calcium activity when subjected to noxious stimuli (**Fig. 5d**).

**Fig. 5.**
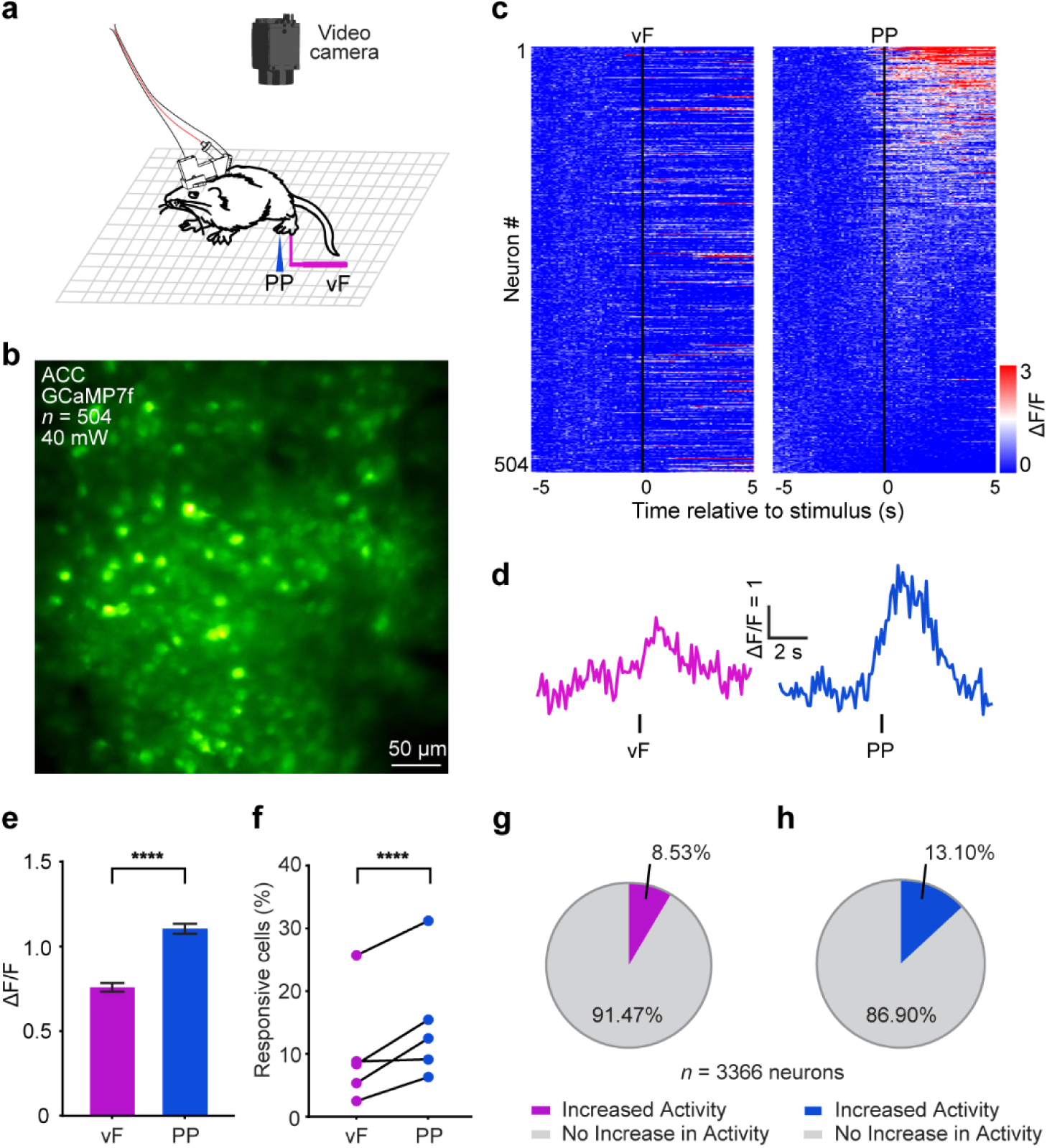
Imaging neuronal populations in ACC under different pain stimulation paradigms. **a**, Schematic diagram for monitoring the response of ACC neurons to noxious (PP, pin prick) or nonnoxious (vF, 0.4 g vF filament) stimuli in freely moving mice. **b**, Image of an example neuronal population in ACC. **c**, Z-scored mean ΔF/F of five trials for 504 neurons recorded from the example mice. Neurons were aligned from high to low calcium responses after PP stimuli. **d**, Representative trace of z-scored mean calcium response of a single neuron during vF and PP. **e**, PP induced a higher calcium response than vF (mean ± s.e.m.); p < 0.0001, paired Student’s t test. **f-h**, More neurons responded to PP than to vF stimuli, p < 0.0001, Fisher’s exact test (**f**); percentages of neurons activated by vF (**g**) and PP (**h**) stimuli (n = 3366 neurons in five mice). Data in **b-d** are from a single representative experiment (n=5 animals, one experiment per animal); data in **e-h** are statistics derived from all five experiments.

We then analyzed the average peak ΔF/F in response to both non-noxious and noxious stimuli, which revealed a significantly higher response to noxious stimuli in comparison to non-noxious stimuli (**Fig. 5e**). Furthermore, more neurons reacted to the noxious stimuli than to the non-noxious stimuli (**Fig. 5f**). Specifically, 13.10% of the neurons recorded responded to the noxious stimuli, whereas 8.53% responded to the non-noxious stimuli (**Figs. 5g-h**). These observations are consistent with previous studies showing increased firing rates in ACC neurons during noxious stimuli [25–26].

Last, we imaged calcium activity of interneurons in ACC upon receiving pin prick stimuli, both before and after chemogenetic inhibition of neural activity in S1 (**Supplementary Figs. 11a-b and Methods**). S1 and ACC are well-known brain regions for processing the sensory and affective components of pain. The S1→ACC pyramidal neuron projection allows sensory pain information to be transmitted to a higher-order cortical center that regulates the affective experience [26]. However, it is not well understood whether ACC interneurons also receive nociceptive information from S1. Using miniBB2p, we found that interneurons in ACC became more sensitive to pin prick after the inactivation of S1(**Supplementary Figs. 11b-c**), suggesting they receive nociceptive information from S1 [26]. This relationship highlights the role of these neurons in processing the sensory and affective aspects of pain.

## Discussion

The Bessel beam has been effectively demonstrated as a promising approach in tabletop 2PM to achieve high-throughput volumetric imaging of neural activity in head-fixed mice across a wide range of neurobiological studies [15–20]. In this work, we found that the unique characteristics of Bessel beams in two-photon imaging—such as extended depth of field, higher lateral resolution, and insensitivity to optical aberrations compared to commonly used Gaussian beams—can facilitate the design of miniature multiphoton microscopes. These advancements not only simplify the design of miniature microscopes but also expand their capability to monitor neural dynamics from a 3D volume at synaptic resolution in freely moving animals, addressing a significant challenge encountered by existing technologies.

The extended depth of field in our miniBB2p system, as in tabletop Bessel-beam 2PM, is essential for high-throughput volumetric imaging but inherently raises the possibility of misinterpreting axially overlapped neurons, particularly in samples with dense labeling. To mitigate this limitation, advanced computational demixing methods, such as constrained nonnegative matrix factorization (CNMF), can be applied to leverage the temporal sparsity of neural activity and separate signals from superimposed neurons [19, 27]. The accuracy of this spatiotemporal signal separation can be further improved by integrating prior structural information acquired from a tabletop Gaussian-beam 2PM, which provides a high-resolution anatomical reference for precise segmentation. Finally, while axially overlapped neurons could lead to noise cross talk, the high sensitivity of advanced genetically encoded calcium indicators renders this effect minimal for resolving individual neurons in calcium imaging [19].

Similar to existing MEMS-based miniature Gaussian-beam 2PMs, miniBB2p achieves an imaging throughput (effective pixel rate) of 2.4 megapixels per second, mainly constrained by the bandwidth of the MEMS scanners. However, miniBB2p delivers a 4–8× improvement in information throughput, enabled by its extended depth of field. Further increases in imaging throughput and FOV would require MEMS scanners with a larger mirror size and wider optical scan angle without compromising the scanning speed. Note that, as the imaging throughput and FOV increases, higher fluorescence excitation and detection efficiency become essential to compensate for the signal loss resulting from reduced pixel dwell times.

The miniBB2p is characterized by its lightweight design, reduced tethering stress, simple optics, and maximized fluorescence collection efficiency. These strengths facilitate long-term stable imaging without impeding the animal behavior, using a safe laser power (40–100 mW) comparable to that used in tabletop Bessel-beam 2PMs (**Supplementary Table 2**) [17]. The high-throughput volumetric imaging capability of miniBB2p enabled us to investigate the neural activities in M2 in response to animal movement, as well as the neuronal responses in ACC under various pain stimulation paradigms, at a large population scale. Higher throughput could be realized by integrating a rapid axial scanning unit into miniBB2p for volumetric imaging across multiple depths [5, 10]. While the large-mode-area fiber used in this work can effectively deliver femtosecond laser pulses over a broad spectral range, a promising feature for imaging indicators with different excitation wavelengths, we noted that the pulse is substantially broadened at high power due to nonlinear effects within the fiber (**Supplementary Fig. 2b**). This shortcoming would compromise performance of the microscope when utilizing a power exceeding 100 mW. Future developments using hollow-core photonic bandgap fiber (HC-920, NKT Photonics) can address this issue and further enhance the sensitivity of miniBB2p in imaging GFP-based indicators [2].

Using readily available off-the-shelf lenses, we developed miniBB2p to be cost-effective without impairing its capability (**Supplementary Table 3**). In addition, the simple optical design allowed us to assemble the lenses in the lab to approach diffraction limit without the need for complex and expensive assembly processes typically required in a lens manufacturing facility. We anticipate that the design’s simplicity and cost-effectiveness will diminish the barriers to a broader application of head-mounted multiphoton microscopy for studying neural dynamics in freely moving animals. Moreover, the optical design methodologies established in this study may pave the way for future advancements in miniature multiphoton microscopes with a larger FOV and higher resolution.

## METHODS

### Microscope setup and system control

A fiber-based femtosecond laser (FemtoFiber ultra 920, Toptica) with a built-in dispersion precompensation unit (-40000 to +1000 fs^2^) was used to produce femtosecond laser pulses at a center wavelength of 920 nm (920±10 nm, 100 fs, 80 MHz, 1.2 W). Using an objective (LMPLFLN20×, Olympus), the laser was coupled into the LMA-12 fiber. Dispersion from the 1-meter fiber was fully compensated with the precompensation parameter set to -26000 fs^2^. In miniBB2p, fiber tip with the custom axicon was aligned with a collimating lens (45-956, Edmund Optics) to generate the ring-shaped beam. The MEMS scanner (A3I12.2-1200AL, Mirrorcle) provided a 1.2-mm diameter mirror with the first resonant frequency of both axes at ∼2.6 kHz and it was used at a 30° incidence angle. After the scan lens and tube lens, which were formed by four identical plano-convex lenses (48-704, Edmund Optics) in a symmetrical configuration, a dichroic mirror (DMR-805SP, LBTEK; 1 mm thick; cut into pieces of 7 × 7 mm^2^ by Sunlight) was used to separate the laser beam and fluorescence. The objective, consisting of two plano-convex lenses (47-850 and 65-300, Edmund Optics), was used to generate the Bessel focus in the specimen. Fluorescence was collected through the objective, dichroic mirror, and filters, and was detected with an onboard miniature, low-cost silicon photomultiplier (3 × 3 mm^2^, S13360-3075PE, Hamamatsu). A reflective filter (FIL-800, 450–650 nm transmission, 6 × 6 mm^2^, 0.2 mm thick, Sunlight) and a colored glass filter (FGB37-A, Thorlabs; 2 mm thick; cut into pieces of 6 × 6 mm^2^ by Sunlight) were placed before the detector to reject the strong infrared laser. The microscope housing was made of carbon-fiber-enhanced plastic (PEEK-CF30) to reduce the weight of the headpiece.

The control signals for x-y scanning were generated from a MEMS controller (USB-SL MZ, Mirrorcle), which was also used to drive the acousto-optic modulator in the laser for power control and to trigger the data acquisition. The scanner was operated in resonance mode along the x-axis at 2304 Hz, scanning 4608 lines every second in bi-directional mode. This operation resulted in an image of 512 × 512 pixels at 9 frames per second. The x-axis scanning range was about √2 times the y-axis scanning range to achieve a nearly square FOV [10]. The silicon photomultiplier was wired by a thin coaxial cable (Ø 260 μm) to the driver circuit (C12332-01, Hamamatsu). Fluorescence signal was amplified by the same circuit and sampled with a digitizer (20 MS/s, M2p.5921-x4, Spectrum Instrumentation). The entire system was controlled by custom software based on the MATLAB (R2022a) platform.

### Micro-axicon design

We customized a micro-axicon at the tip of the LMA-12 fiber (Comcore, China). Specifically, the tip of the photonic crystal fiber was collapsed and polished to form an axicon lens (apex angle = 120°). The laser from the photonic crystal region in the fiber was spread out at the collapsed region (∼1 mm length) and transformed into a diverged ring beam by the axicon (**Supplementary Fig. 2c**). After the collimating lens, a focused ring beam was generated at the back focal plane of the scan lens, where the MEMS mirror was located. The maximum diameter of the ring beam that the mirror can accommodate is 1 mm, corresponding to a maximum excitation NA of 0.3. In this work, the diameter of the ring beam was measured to be 0.85 mm and it resulted in an excitation NA of 0.26.

### Resolution and FOV calibration

We ascertained the resolution of miniBB2p by imaging 500-nm fluorescent beads (F8813, Thermo Fisher Scientific), with a pixel size of 0.1 μm to measure lateral resolution and an axial step size of 6 μm to measure axial resolution. The resolutions were evaluated as the full width at half maximum of the point spread functions (**Supplementary Fig. 3**). We calibrated the FOV of miniBB2p by imaging a 50-μm grid standard (RTR1Ee-P, LBTEK). Field distortion due to optical aberrations (**Fig. 1d**) and MEMS scan was corrected with the software *DistortionCleaner* (**Supplementary Figs. 4e-f**) [10]. All data in this study was displayed after distortion correction.

### Tabletop Gaussian-beam 2PM

A tabletop Gaussian-beam 2PM (AX R MP, Nikon) with a water immersion objective (CFI Plan Fluor 10× W, NA 0.3, Nikon) was used to evaluate the performance of the miniBB2p. Lateral and axial resolution of the tabletop 2PM measured from 500-nm fluorescent beads were 1.34 ± 0.02 μm and 22.90 ± 0.47 μm (mean ± s.d., n = 5).

### Animals

C57BL/6J male wild-type mice (Jackson Laboratory stock No. 000664) were obtained and housed at the Laboratory Animal Resource Center (LARC) within the Chinese Institute for Brain Research (CIBR). Thy1-GFP-M and VGAT-Cre transgenic mice were generously donated by Wenzhi Sun at CIBR. All mice were acquired at 5 weeks of age and allowed an average of 2 weeks to acclimate to their new environment before the experiments began. All experimental procedures were approved by the Institutional Animal Care and Use Committee of CIBR, in accordance with the governmental regulations of China.

### Viral construction and packaging

Recombinant adeno-associated virus (rAAV) vectors were serotyped with AAV9 coat proteins and packaged at BrainCase viral vector manufacturing facilities. The viral titers were approximately 3.3 × 10^12^ particles per milliliter for rAAV2/9-CaMKIIα-hM4D(Gi)-mCherry and 5.2 ×10^12^ particles per milliliter for AAV-CaMKIIα-jGCaMP7f. Sparse expression was achieved by injecting a mixture of diluted AAV2/9-hSyn-Cre particles (original titer: ∼1.3 × 10^13^ particles per milliliter, diluted 5,000-fold in sterilized phosphate-buffered saline) and high-titer, Cre-dependent GCaMP8s virus (rAAV2/9-hSyn-DIO-jGCaMP8s, 5.2 × 10^13^ infectious units per milliliter). Viral aliquots were stored in a light-protected freezer until use.

### Surgical procedures

Mice were anesthetized with 1.5%-2% isoflurane, and viral vectors were delivered to the ACC, M2, or S1 regions of the brain. A fitted plunger controlled by a hydraulic manipulator (Beijing Xinglin Life Tech.) was inserted into a glass pipette to load and slowly inject 50 nL of viral vectors at a rate of 2 nL/5 s. Injections were performed bilaterally in the ACC at anteroposterior (AP) +1.3 mm, mediolateral (ML) ±0.3 mm, and dorsoventral (DV) -0.6 mm; bilaterally in S1 at AP -0.59 mm, ML ±1.5 mm, and DV -0.4 mm; and unilaterally in M2 at AP 0.6 mm, ML ±0.75 mm, and DV -0.5 mm. Following the injection, the microinjection needles were left in place for 10 minutes to allow viral diffusion and minimize the spread of viral particles along the injection tract. The pipette was then left in position for an additional 5 minutes before being slowly raised out of the brain.

After 3 weeks of expression, the mice were anesthetized with 1.5% isoflurane for cranial window implantation. Analgesic compound lidocaine cream was applied to the skin 10 minutes before the surgical incision, and ophthalmic ointment was used to prevent eye drying. The body temperature was maintained at 36°C using a feedback-controlled heating pad (FHC). A craniotomy with a diameter of 4.3 mm was performed over either the ACC or M2, leaving the dura intact. A glass coverslip of matching diameter was placed over the cranial window, and the gap between the glass window and the skull was sealed with Vetbond (3M, USA). After that, the isoflurane concentration was reduced to 0.5%-1%. The miniBB2p, with a custom titanium baseplate attached, was adjusted relative to the glass window to find the target plane for free-moving neural imaging. The baseplate was then secured with dental acrylic, and a custom cover was placed on the baseplate, fixed with screws, to protect the cranial window when not in use. Before surgery and testing, the mice were handled and pre-trained in the recording environment until they were acclimated and foraged freely within the box.

### Mice handling and pretraining

Mice were habituated to the experimental environment for 3 days prior to locomotion test or free-moving imaging. During the same period, they also underwent daily handling sessions to acclimate to head-mounted procedures. Handling included gentle headcap pressing, back stroking, and holding the headcap firmly for approximately 10 seconds. Each session lasted 5–10 minutes and was designed to reduce stress and facilitate later dummy or real miniBB2p mounting in the awake state. All handling and behavioral procedures were conducted under simulated night time conditions. For locomotion test, awake mice were fitted with a dummy miniBB2p (or not, for controls) and allowed to habituate to the experimental environment for 30 minutes. They were then gently transferred to the open field arena for a 5-minute exploratory adaptation, followed by a 30-minute free-movement recording session (*EthoVision XT 16*, 10 Hz sampling). For free-moving imaging, the real miniBB2p was mounted while the mice were awake, followed by a habituation period in the imaging chamber placed over a mesh table. Once the animal exhibited stable and naturalistic movements, imaging recordings were initiated.

### Imaging procedures

At the beginning of each imaging session, the mice were head-fixed and the miniBB2p was mounted on the baseplate. Behaviors of freely moving mice were recorded with a camera (MV-CS013-60GN, HIKROBOT) during entire experiment. Initially, spontaneous neural activity was recorded while the mice habituated to the testing environment, freely exploring the 30 cm × 20 cm × 20 cm recording chamber without any experimenter-delivered sensory stimuli.

For noxious stimulation, the plantar surface of the hind paw contralateral to the brain recording site was pricked with a 27-gauge needle (pin prick) while the mice were freely moving. The stimulation ceased upon the withdrawal of the paw. Non-noxious stimulation was delivered to the same hind paw using a 0.4 g von Frey filament, applied continuously for 3 seconds or until paw withdrawal. In most cases, there was no withdrawal response to the von Frey filament. Each recording session included approximately 10 trials with variable inter-trial intervals of around 30 seconds to prevent sensitization. Throughout the sessions, no signs of behavioral sensitization or physical damage to the paws were observed.

### Motion Correction and image analysis

To eliminate motion artifacts in the calcium imaging data from each session, piecewise non-rigid motion correction was applied using *NoRMCorre* [28] or *Flow-Registration* [29] (used in **Supplementary Fig. 7**). The motion-corrected 9 Hz calcium activity videos were saved as .TIFF files and subsequently used for cell identification.

For C57BL/6J mice injected with AAV-CaMKIIα-jGCaMP7f, calcium signals were extracted using *Suite2P*, which constrains the instances of calcium activity for each neuron [30]. The analysis produced three traces for each identified region of interest (ROI), corresponding to putative cell somata: the raw fluorescence signal F_cell_(t), the neuropil fluorescence signal F_np_(t), and the deconvolved calcium activity E(t). F_cell_(t) and F_np_(t) represent the weighted averages of activity within the ROI and the surrounding neuropil, respectively, excluding pixels overlapping with other ROIs. To correct for neuropil contamination, the neuropil signal was subtracted from the raw calcium signal using a fixed coefficient [31], yielding the corrected signal: F_corr_(t) = F_cell_(t) - 0.7 × F_np_(t). To mitigate the effects of slow fluctuations such as photobleaching on F_corr_, a running baseline F_0_(t) was extracted. The calcium activity was then expressed as the fractional change in fluorescence relative to the baseline ΔF/F, calculated as ΔF/F = (F_corr_(t) - F_0_(t)) / F_0_(t).

For VGAT-Cre transgenic mice injected with rAAV2/9-hSyn-DIO-jGCaMP8s, the calcium imaging data from VGAT^+^ cells were analyzed using the commercial image-analysis platform AIVIA 11.0 (DRVision Technologies LLC, Bellevue, WA) in conjunction with the *Cellpose* plugin. The segmentations from *Cellpose* were converted into masks for individual neurons, and the averaged fluorescent signal within each mask was used to calculate calcium transients.

### Locomotion test

Custom-made dummy miniscopes (2.6 g) were utilized to assess the impact of miniBB2p on the behavior of mice during imaging. These dummies replicated the shape and weight distribution of the actual miniBB2p and were mounted onto implanted headbars, accompanied by one fiber and two cables. Twelve adult C57BL/6 mice (n=6, 4 M/2 F, real implants with ACC cranial windows; n=6, 3 M/3 F, sham implants with headbar-only) were co-housed (2-3 per cage) under a 12 h reversed light/dark cycle. All mice underwent two counterbalanced trial phases: a control phase involving open-field exploration in a white acrylic arena (97.6 cm × 45 cm × 38 cm) without attachments, and a dummy phase following the same procedure but with a mounted dummy miniscope. Each subject completed one trial type per 24 h cycle. Locomotor parameters including total distance traveled, mean/median velocity, central zone entries, and distance distribution (central vs corner zones) were quantified using the *EthoVision XT 16* tracking software.

### Brain slice preparation

Mice were deeply anesthetized with isoflurane and then transcardially perfused with ice-cold phosphate-buffered saline (PBS) followed by paraformaldehyde (PFA). After extraction, the brains were fixed overnight in PFA. The fixed brains were then sectioned into 85 µm-thick slices using a vibratome (Leica VT1200S). These brain slices were mounted onto slides using a mounting medium containing DAPI (Solarbio, China) to preserve the fluorescent signal for long-term storage.

### Calcium response analyses

For the pain stimulation experiments, trial-averaged calcium fluorescence traces were generated for individual neurons by z-scoring each trial from -5 to 5 seconds, with 0 indicating the time of stimulation. Z-scoring involved subtracting the mean and dividing by the standard deviation of a baseline period from -5 to -3 seconds prior to pinprick stimulation. The z-scored trials were then summed and divided by the total number of trials, resulting in a single trial-averaged calcium fluorescence trace for each neuron.

To obtain a session-level analysis, trial-averaged calcium fluorescence traces from individual neurons were combined and averaged across all neurons, producing a single representative trace of neuronal activity. The peak calcium fluorescence for each neuron was identified within the post-stimulation window of 0 to 5 seconds. Each trial was z-scored as described, and then binned into 1-second intervals. A 2-second moving window was applied to determine the maximum average calcium fluorescence for each trial. The peak values from all trials were then averaged to yield a singular value representing the peak post-stimulus activity of each neuron in a given session.

To evaluate the effect of running on neural activity in M2, we binarized the behavior of the mice into running/resting states and compared their mean ΔF/F values for each neuron. The running modulation index (MI) is calculated as (ΔF/F_run_−ΔF/F_rest_)/(ΔF/F_run_+ΔF/F_rest_). A running onset was detected when a period of at least 400 ms of resting was followed by at least 400 ms of running [24]. For the investigation of neuronal response to running onsets, we inspected the ΔF/F responses with onsets that met two criteria simultaneously: (1) the onset had to be at least 10 seconds away from previous onset, and (2) the running episode had to last at least 1 second. This operation ensured that only well-separated and sustained running episodes were analyzed, minimizing the influence of transient or ambiguous movement signals.

### Chemogenetics

Clozapine N-oxide (CNO; APExBIO, China) was prepared as a suspension in 0.9% saline, achieving a final concentration of 0.2 mg/mL. The solution was administered via intraperitoneal injection at a dose of 1 mg/kg. To activate the hM4D(Gi) DREADD, imaging was conducted 30 minutes following the systemic (intraperitoneal) injection of CNO.

### Behavior tracking

The raw tracking video recorded during free-moving imaging was initially analyzed using *DeepLabCut* (*DLC*) [32], which provided the positional data for specific body parts of the animal across each frame. Three key body parts were selected for analysis: the head, the body center, and the tail base. The *DLC* model was trained on a minimum of 20 frames, following the default training protocol, and underwent 30 thousand iterations. This training resulted in a final loss of approximately 0.003, with a training error of 1.48 pixels and a test error of 3.33 pixels. The animal’s position was recorded in x-y coordinates as x(t) and y(t). To smooth the positional traces, a low-pass filter was applied to both x(t) and y(t). The instantaneous movement speed was subsequently calculated by dividing the displacement of the head between consecutive frames by the time interval (1/frame rate), yielding a speed in centimeters per second (cm/s). Two-photon imaging was synchronized with the animal tracking camera in hardware, enabling precise temporal alignment between neuronal activity and behavioral tracking data. Calcium fluorescence traces were captured with the same number of frames as the tracking signals, ensuring that time points were pre-aligned across both datasets.

### Multi-day registration and cross-validation with a Gaussian-beam 2PM

We first recorded neuronal activity in vivo using miniBB2p and later acquired the high-resolution 3D morphological data from the same imaging volume using a tabletop Gaussian-beam 2PM (objective: CFI Apo NIR 40×, NA 0.8, Nikon) on the same day. This paired acquisition protocol was repeated for the same imaging volume across multiple recording days.

Cell registration across multiple recording days was performed using the *ROIMatchPub* MATLAB package (https://github.com/ransona/ROIMatchPub) after functional segmentation with *Suite2p*. The registration pipeline consisted of three main steps: (1) manual labeling of reference cells and landmarks across multi-day recordings, (2) automatic detection of potential cell matches based on a 50% overlap ratio threshold [10], and (3) manual validation of matched cell candidates through morphological analysis to eliminate false positives. The registration reliability was further assessed through cross-validation between the 2D volumetric projections from miniBB2p with high-resolution 3D images from a tabletop Gaussian-beam 2PM. The 3D images were first segmented using Cellpose3 [33], with subsequent refinement through AI-trainable segmentation in Imaris (v10.2) and manual verification based on morphological criteria. After that, the miniBB2p cell registrations generated by *ROIMatchPub* were systematically validated against their corresponding 3D segmented counterparts.

### Statistical tests

For imaging data, paired neuronal recordings from a single session were compared using a paired Student’s t-test, while unpaired data from separate sessions were analyzed using an unpaired Student’s t-test. Changes in neuron populations before and after drug administration were assessed with Fisher’s exact test. To determine pain-responsive neurons, a one-tailed Wilcoxon rank sum test was conducted on pre- and post-stimulus values for all peripheral stimulations during pain stimulus sessions, with significance set at p < 0.025. A similar one-tailed Wilcoxon rank sum test was employed to identify the neurons that responded to running onsets by comparing calcium activity during resting and running periods. One-way repeated measures ANOVA with Tukey post hoc tests were used to assess the effects of photodamage in chronic recordings.

### Data processing

Images were visualized and processed using the open-source software ImageJ. Unless stated otherwise, all images presented here were unprocessed raw images without smoothing, denoising, and deconvolution. “Green hot” lookup table in ImageJ was used for all miniBB2p captured images. Cross-section profiles, statistical graphs, and other coordinate graphs were drawn with Origin 2022 or MATLAB (R2022b). All neural activities and behaviors were collected at 9 and 27 fps, respectively. The supplementary videos were made at 25 fps using Adobe Premiere Pro 2024.

## Data availability

The optical design files, technical drawings, assembly protocol of miniBB2p are available in **Supplementary Data** (https://github.com/WuLab-CIBR/miniBB2p). The data supporting the findings are provided in the main text, supplementary figures and videos. The Source Data are provided with this paper. All data underlying this study are available from the corresponding author upon reasonable request.

## Code availability

The code for hardware control of miniBB2p are available in **Supplementary Data** (https://github.com/WuLab-CIBR/miniBB2p).

## Acknowledgments

This work was supported by the Innovation Fund for Medical Sciences from Chinese Academy of Medical Sciences (J. Wu), the startup funds from CIBR (J. Wu), the Beijing Municipal Science & Technology Commission (Z220009 to J. Wu), the National Natural Science Foundation of China (52205550 to L. Qian and 82401719 to Y. Liu). The authors thank the Behavior Analysis Center, Imaging Core, Instrumentation Core, Vector Core and LARC in CIBR for their support, and thank W. Z. Sun and M. M. Luo for constructive comments on the free-moving studies.

## Contributions

J. Wu conceived the project and designed the optics; L. Qian designed the mechanics and electronics, L. Qian constructed the microscope; Y. Liu designed all animal research; Y. Liu, L. Qian, and Y. Chen collected data; Y. Liu, L. Qian, and Y. Chen analyzed data; J. Wu supervised research; J. Wu, Y. Liu, L. Qian, and Y. Chen wrote the manuscript with inputs from all authors.

---

**Supplementary Fig. 1.**
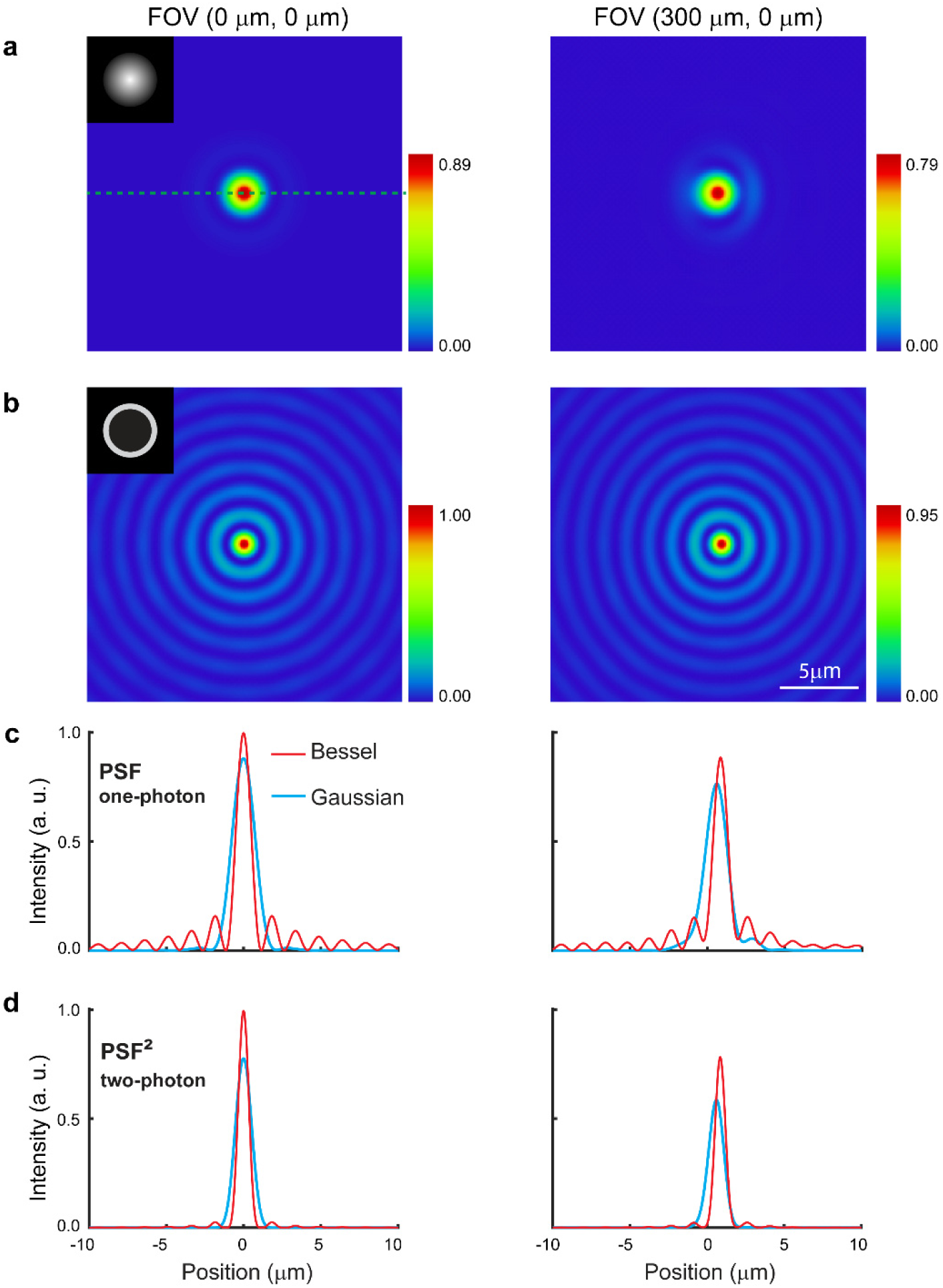
Simulation of miniBB2p with Gaussian and Bessel beam excitation. **a**, Gaussian focus at the focal plane; inset, beam diameter at the stop is 1 mm (NA=0.3). **b**, Bessel focus at the focal plane; inset, annular aperture at the stop, outer and inner diameters are 1 mm and 0.98 mm (NA=0.3). **c**-**d**, Lateral point spread function (PSF) with Gaussian and Bessel beam excitation, **c**, one-photon; **d**, two-photon. Maximum intensity of an unaberrated PSF is 1. Left and right panels compare their performance at the center and edge of the FOV.

**Supplementary Fig. 2.**
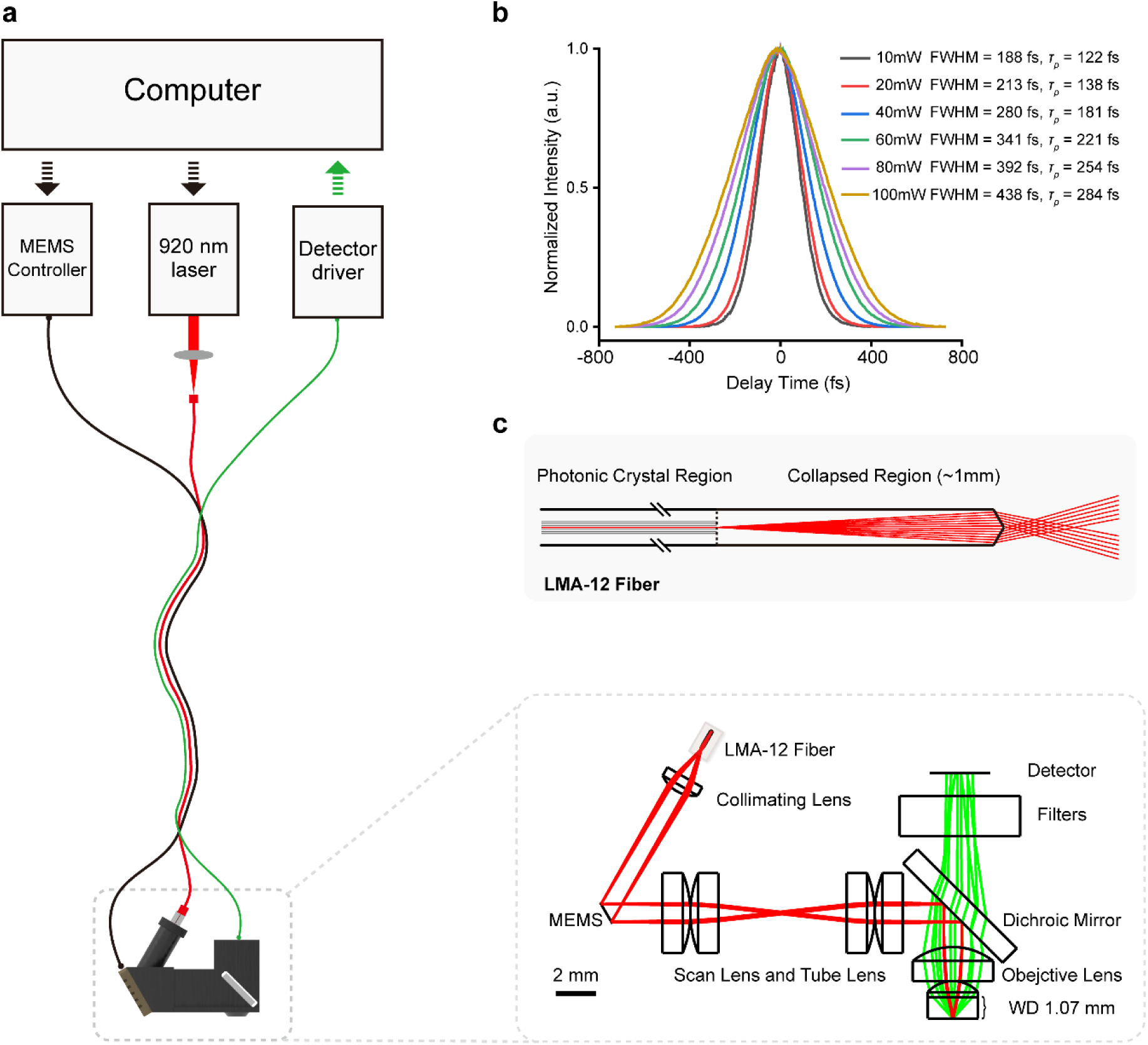
Design of miniBB2p and pulse delivery test. **a**, Overview of miniBB2p; inset, optical layout of miniBB2p headpiece; WD, working distance. **b**, Autocorrelation traces from an autocorrelator (FR-103HP, Femtochrome) showing the widths of 920-nm laser pulses after passing through a 1-meter LMA-12 fiber at different powers. The pulse width (*τ_p_*) is calculated as ∼0.65 times the width of the autocorrelation signal, assuming the pulse is shaped like a sech² function, which is typical in mode-locked lasers. FWHM, full width at half maximum. **c**, Schematic of the photonic crystal fiber LMA-12 with a custom axicon at the tip.

**Supplementary Fig. 3.**
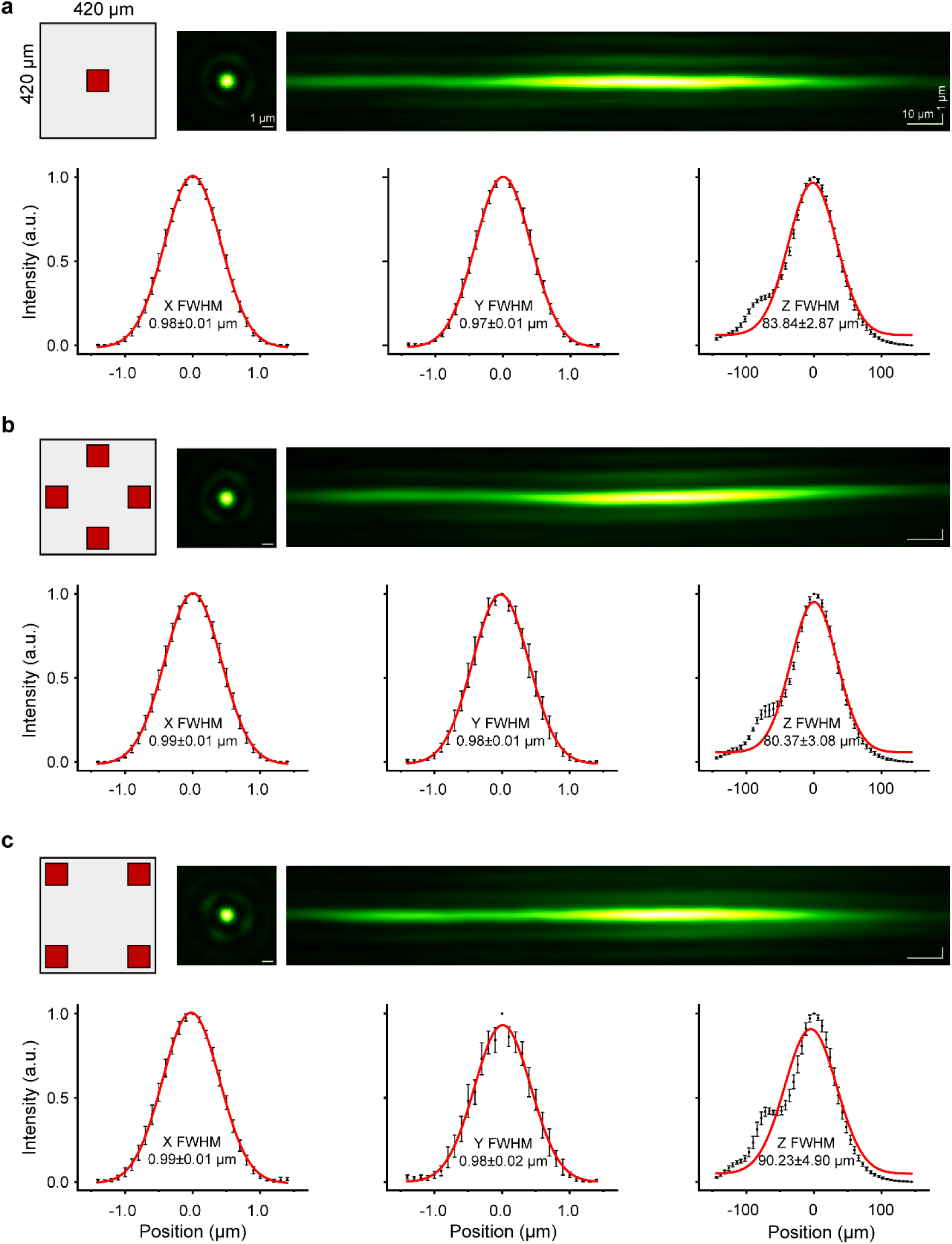
Lateral and axial resolution in miniBB2p. PSF at the center (**a**); edge (**b**), and corner (**c**) of the FOV. Top: illustration of the positions for measuring resolution, representative image and side projection of a 500-nm fluorescent bead. Bottom: PSF profiles; the resolutions were evaluated as the FWHM of the PSFs (mean ± s.d., n = 10).

**Supplementary Fig. 4.**
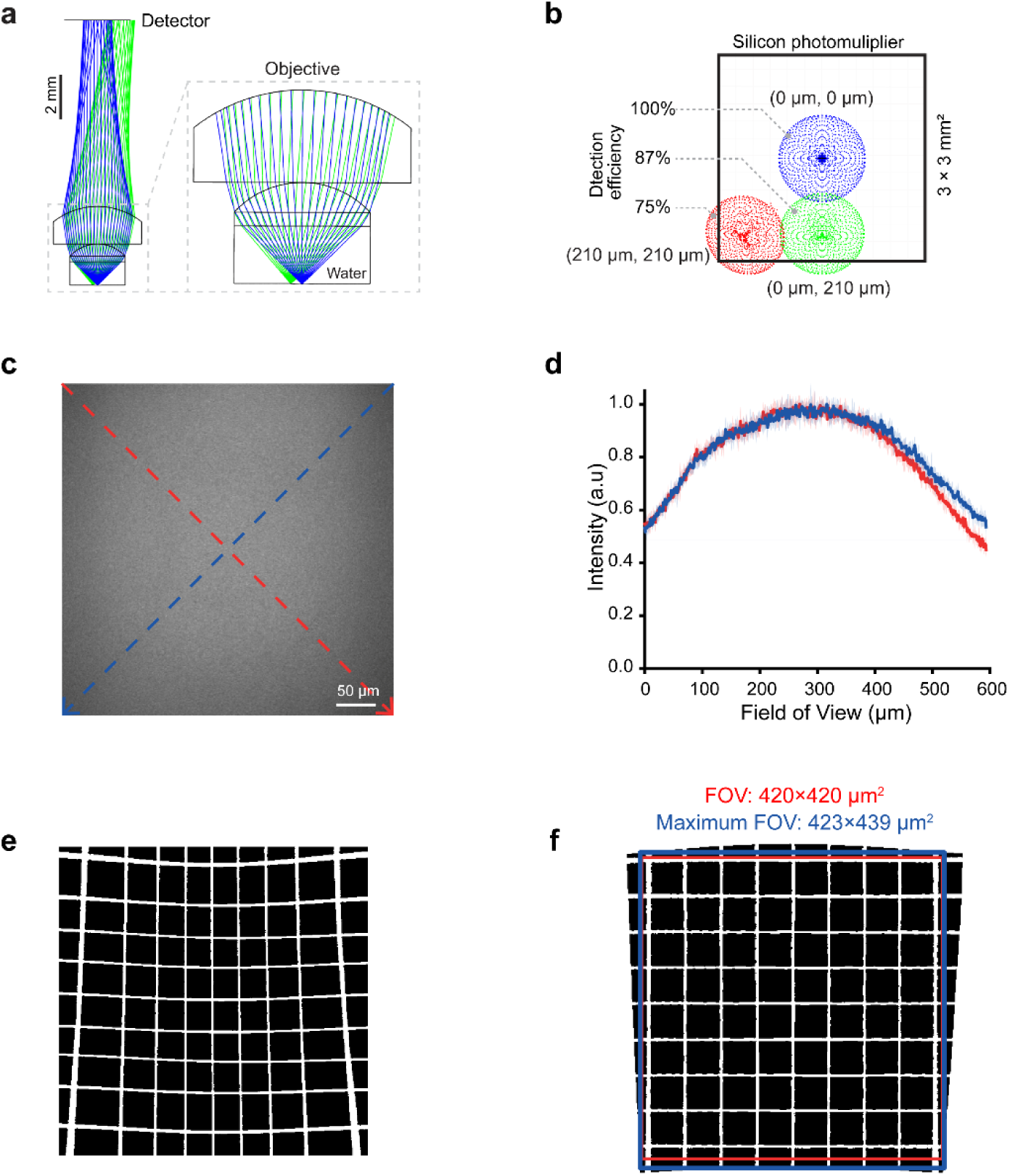
Fluorescence collection and distortion correction in miniBB2p. **a**, Ray path from the focal plane to the detector; zoom-in view shows the ray path in the objective; fluorescence collection NA is 0.95; dichroic mirror and filters are omitted for simplicity. Blue and green rays illustrate fluorescence from the center and edge of the FOV. **b**, Fluorescence from different locations detected by the silicon photomultiplier. The simple optics in miniBB2p enabled highest detection efficiency in the center and an acceptable detection efficiency at the edge (87%) and corner (75%) of the FOV. **c**, Field uniformity tested with a fluorescent plastic slide (92001, Chroma). **d**, Normalized fluorescence intensity across the diagonal lines in **c** (average from ten images, shaded areas show standard deviation). **e-f**, Bright field image of a 50-μm grid standard before (**e**) and after (**f**) distortion correction (see **Resolution and FOV calibration** in **Methods**); image was captured without the filters installed in miniBB2p.

**Supplementary Fig. 5.**
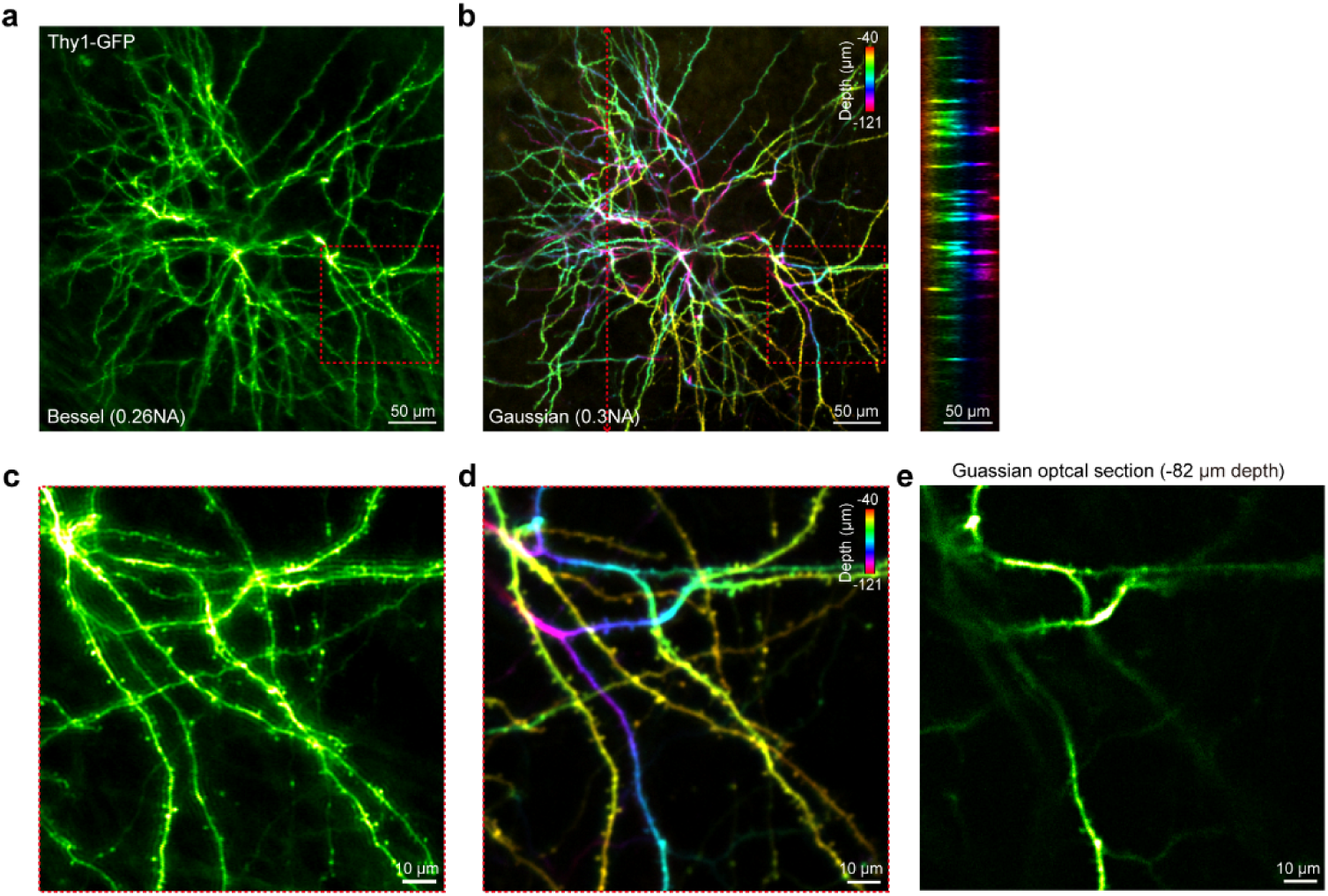
Imaging of dendrites and spines in the head-fixed awake Thy1-GFP mouse brain. **a**, Image of a ∼80-μm thick volume in ACC by miniBB2p. **b**, Lateral projection of an image stack at 3-μm step collected by a tabletop Gaussian-beam 2PM from the same volume, with structures colored by depth (left); cross section view across the dashed line (right). **c-d**, Zoom-in imaging of the boxed areas in **a-b**. **e**, Optical section at the middle-depth in **d.** Data are from a single representative experiment (n=3 animals, one experiment per animal).

**Supplementary Fig. 6.**
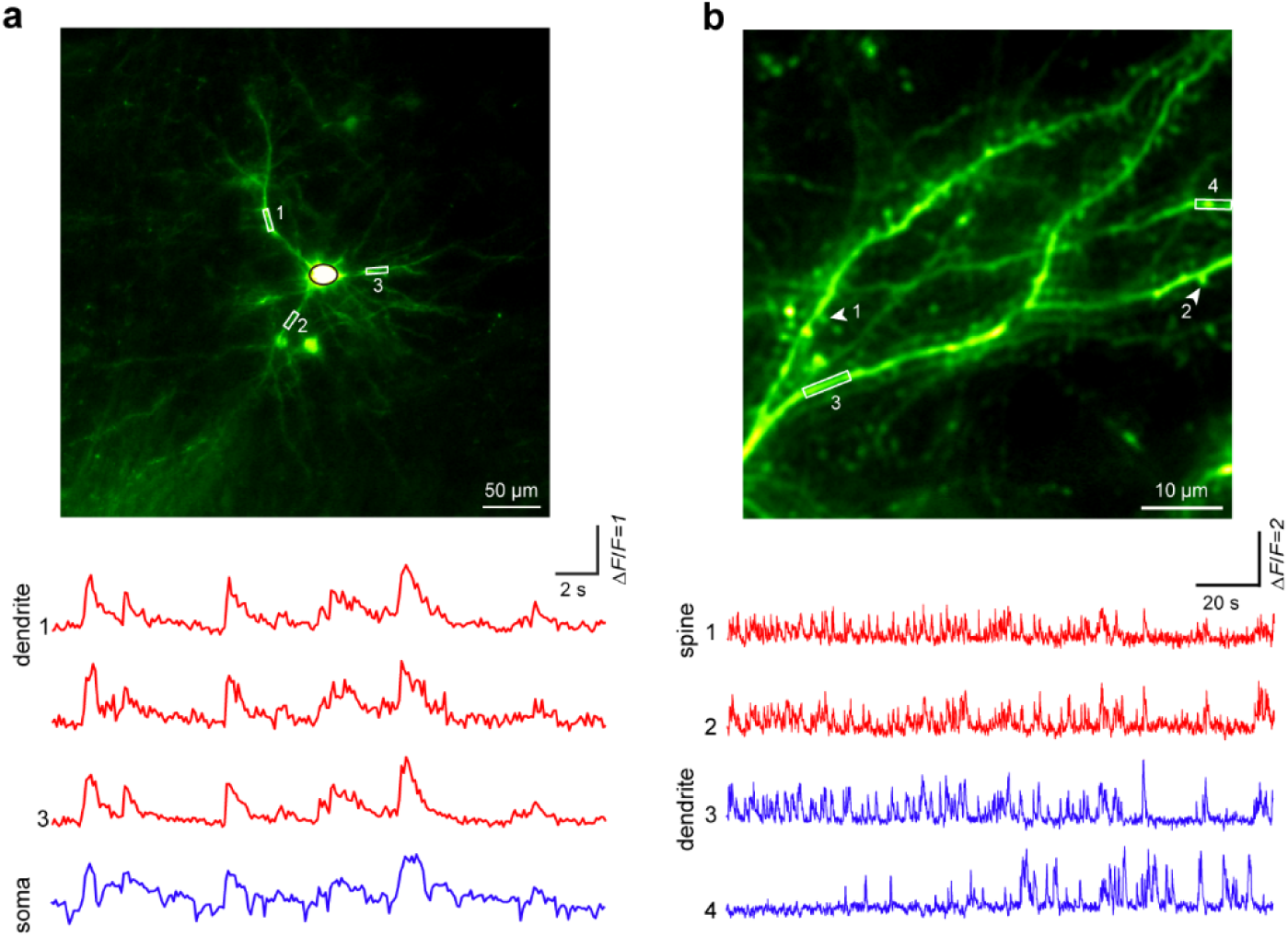
Imaging of the activities of dendrites and spines in ACC expressing GCaMP8s in head-fixed awake mice. **a**, FOV, 420 × 420 µm^2^. **b**, FOV, 60 × 60 µm^2^. Data in **a**-**b** are from a separate representative experiment (n=6 animals, one experiment per animal)

**Supplementary Fig. 7.**
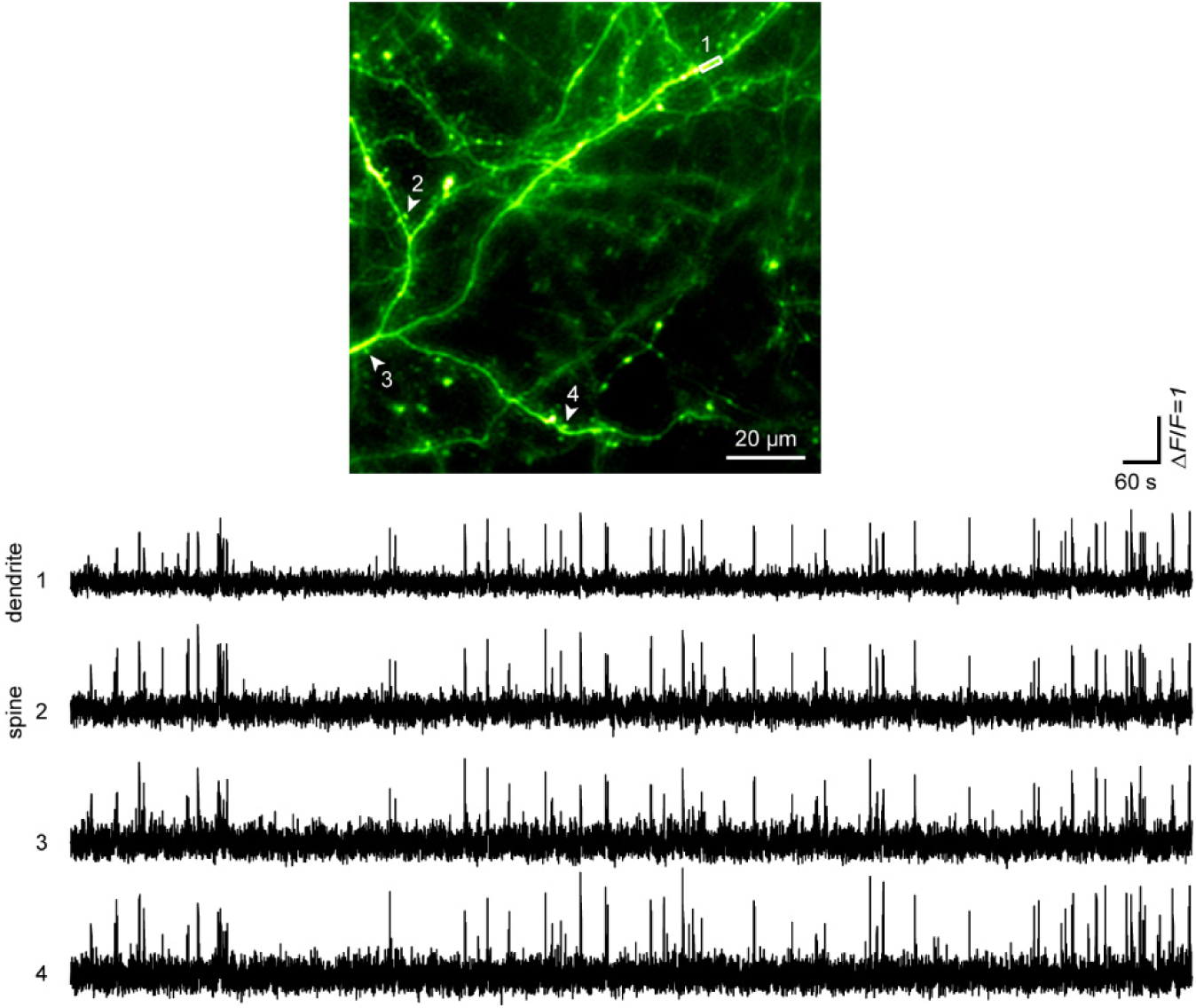
Imaging of the activities of dendrites and spines in ACC expressing GCaMP8s in a free-moving mouse over a 30-minute period. Image of a ∼80-μm thick volume in ACC by miniBB2p (top) and representative calcium traces of dendrite and spines (bottom). Data represent one of three experiments in one animal.

**Supplementary Fig. 8.**
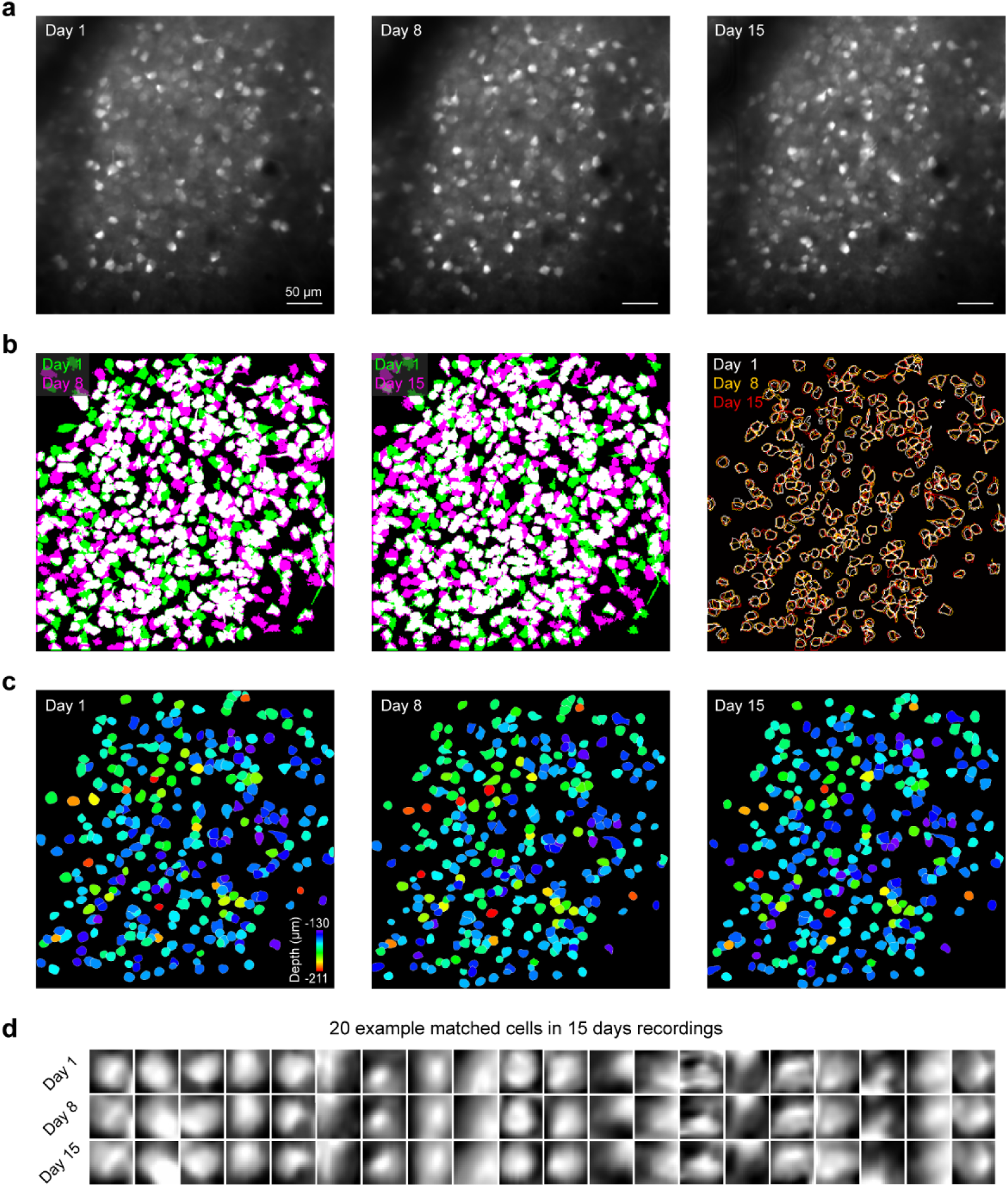
Stability of miniBB2p imaging. **a**, Recording the same volume in ACC across multiple days. **b**, Cell registration across sessions: Day 1 cells (green), Day 8 or Day15 cells (pink), overlapping regions (white). Total cell counts: Day 1 (572), Day 8 (628), Day 15 (670). 65% of Day 1 cells remained detectable after 14 days (372 matched to Day 8; 370 to Day 15), with 53% (301 cells) persisting across all three sessions. **c**, Spatial distribution of the 301 cells persisting across all three sessions, reconstructed from a tabletop Gaussian-beam 2PM 3D imaging data (**Methods**). **d**, 20 example cells identified across all recordings, showing similar cell morphology. Data are from a single representative experiment (n=2 animals, one experiment per animal).

**Supplementary Fig. 9.**
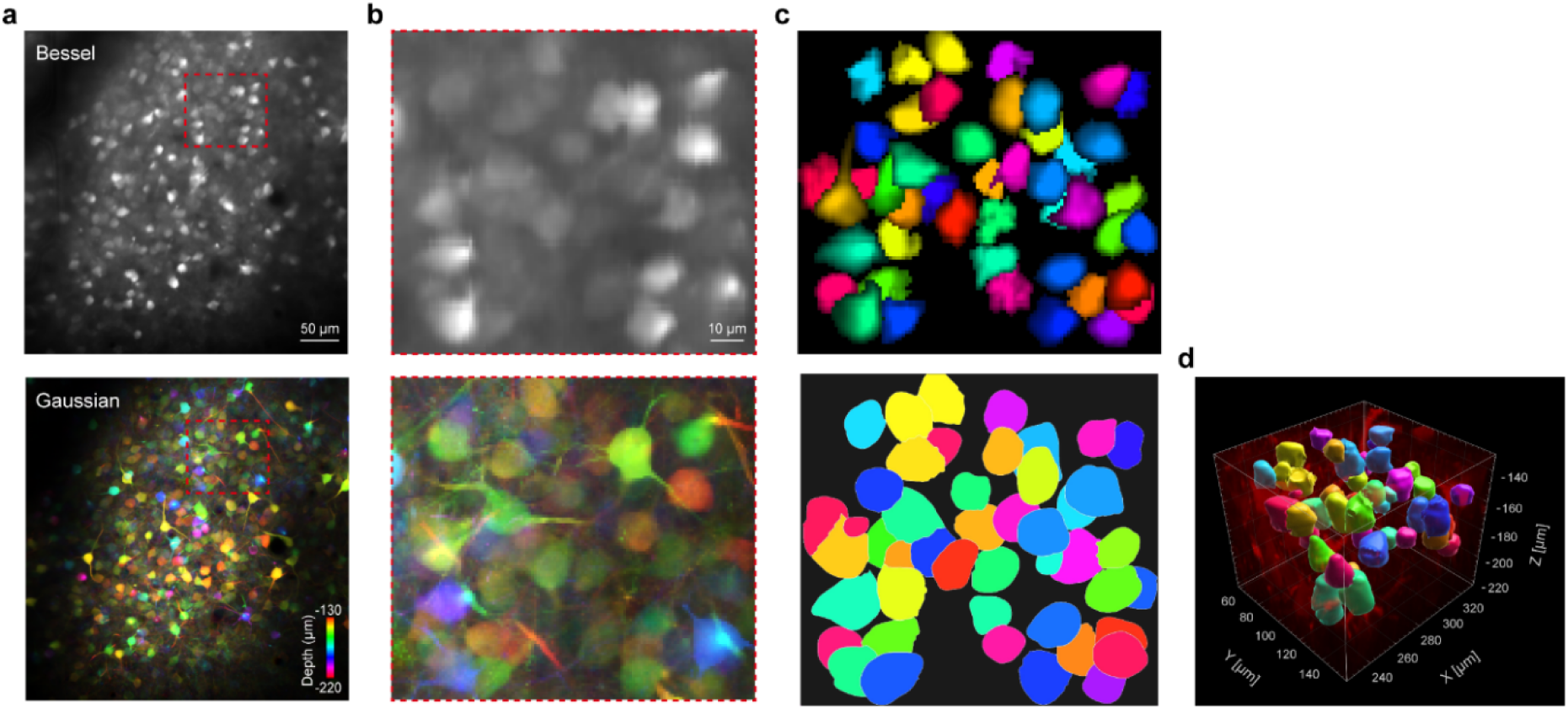
Validation of cellular registration in miniBB2p with a tabletop Gaussian-beam 2PM. **a**, Imaging of the same volume using miniBB2p (top) and tabletop Gaussian-beam 2PM at an axial interval of 3-μm, with structures colored by depth (bottom) (Day 15 data in **Supplementary** Fig. 8). **b**, Zoom-in view of the boxed areas in **a**. Note that weak signals (e.g., from dendrites and spines) in miniBB2p become obscured by stronger somatic signals due to axial signal integration, which is an inherent limitation in Bessel-beam 2PMs. **c**, Cells identified by *Suite2p* from the miniBB2p imaging data (top); cells identified by *Cellpose3* from the Gaussian-beam imaging data (bottom). Cells are randomly colored for visualization. **d**, 3D spatial distribution of the cells in the Gaussian-beam imaging data.

**Supplementary Fig. 10.**
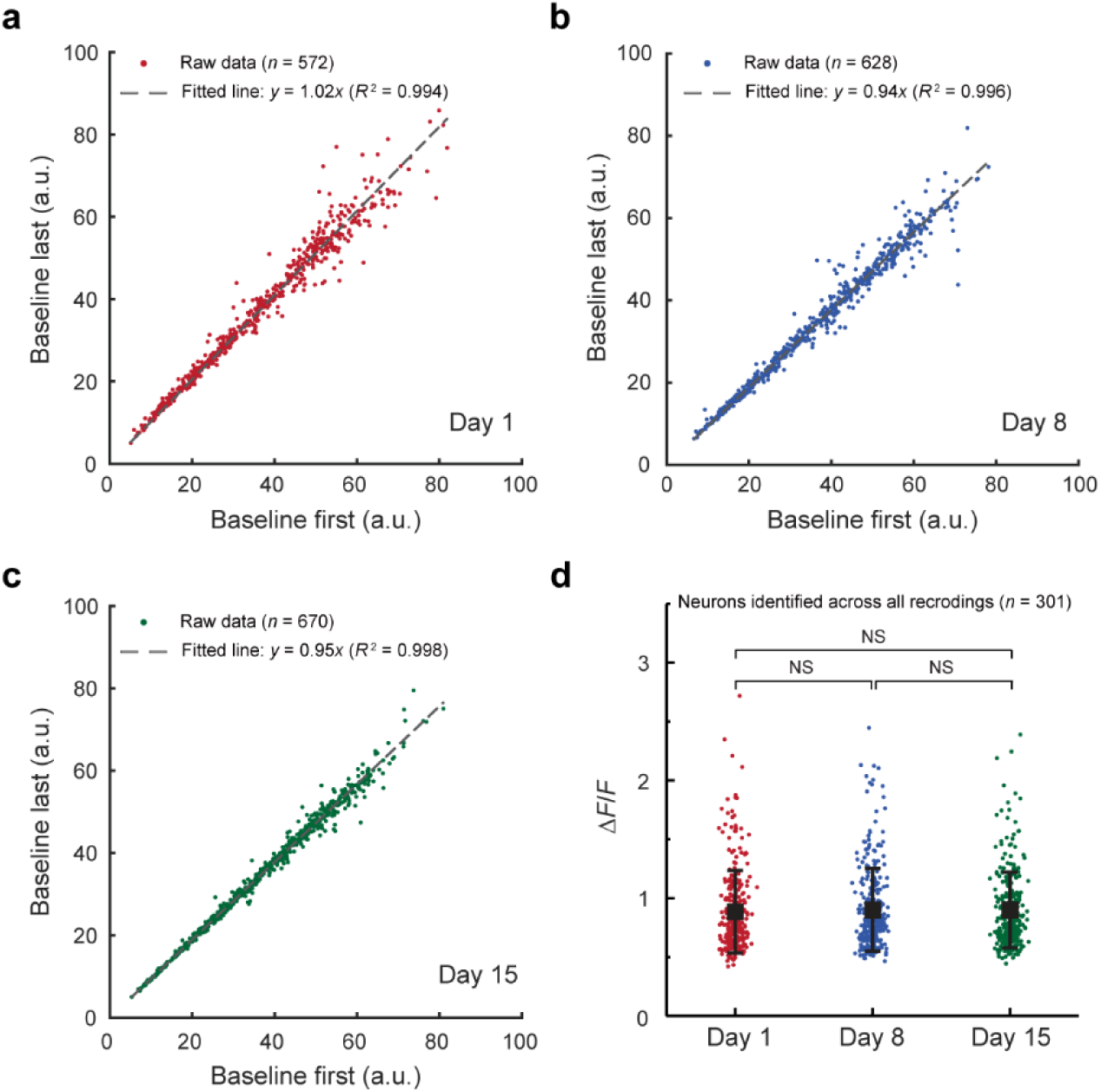
Minimal photobleaching and photodamage in miniBB2p. **a-c**, Neuronal soma fluorescence baselines (20th percentile intensity values) during the first and last 10% of the 30-minute recording period (raw data in **Supplementary** Fig. 7). **d**, Average peak *ΔF/F* of calcium transients measured in the same neuronal populations (Day 1, 0.89 ± 0.35; Day 8, 0.90 ± 0.35; Day 15, 0.90 ± 0.32; mean ± s.d.). No significant differences were observed across the three recordings over two weeks (One-way repeated measures ANOVA with Tukey post hoc tests, Day 1 versus Day 8: p = 0.747, Day 8 versus Day 15: p = 0.998, Day 1 versus Day 15: p = 0.781).

**Supplementary Fig. 11.**
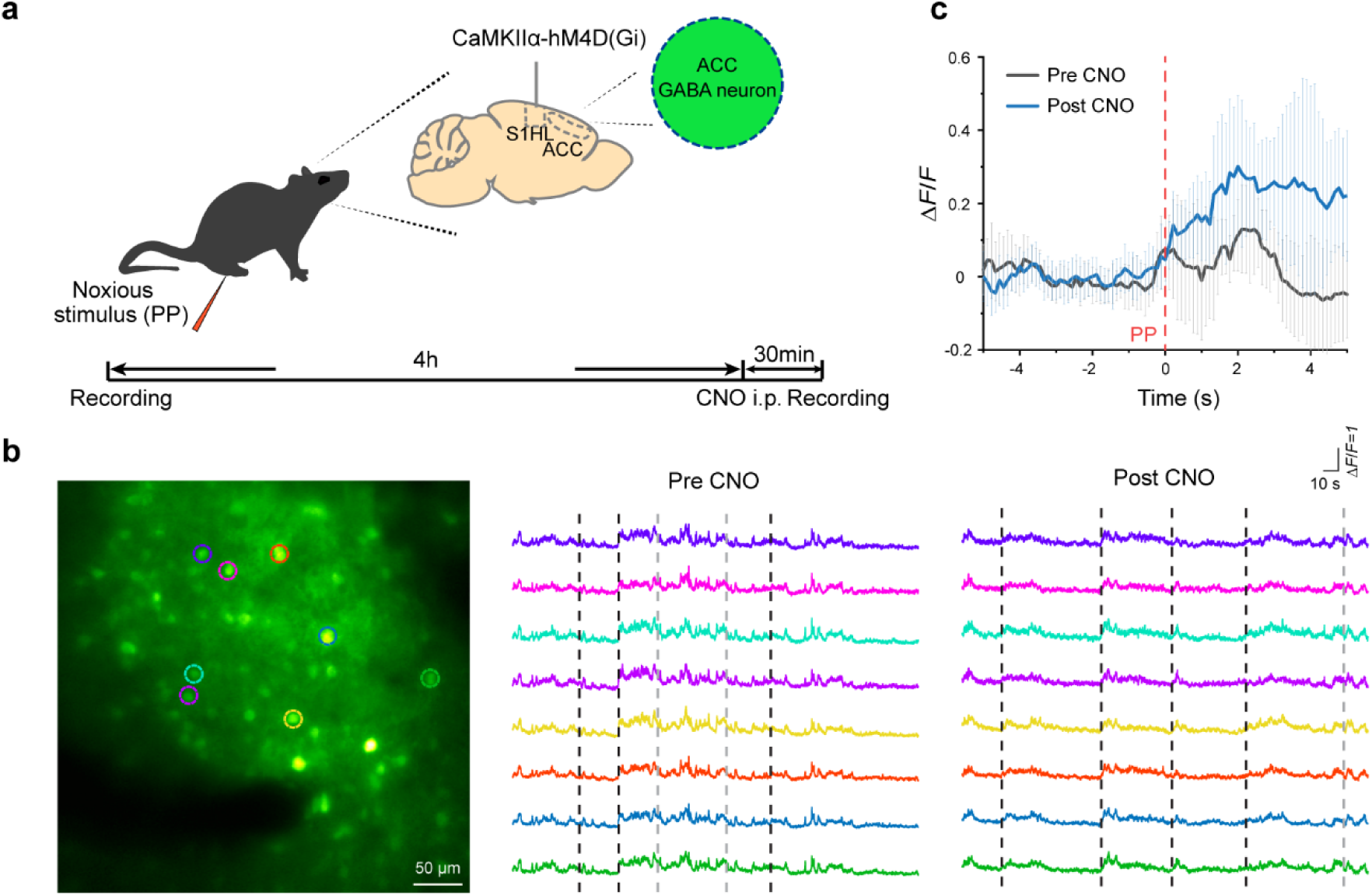
Inhibition of S1 inputs enhance the nociceptive response in ACC. **a,** Schematic of the experiment. **b**, Imaging of ACC using miniBB2p (left), example ΔF/F traces of color-outlined neurons before (middle) and after (right) CNO injection; dashed lines indicate pin prick timings. **c**, Mean calcium responses (± s.d.) of eight neurons color-outlined in **b** during pin prick. Traces with unstable baselines before pin prick (grey dashed lines in **b**) were excluded. Data are from a single representative experiment (n=3 animals, one experiment per animal).

**Supplementary Table 1:** Optical performance in the state-of-the-art miniature multiphoton microscopes.

| Reference | Klioutchnikov, A. et al.<br>*Nat Methods* (2020) | Klioutchnikov, A. et al.<br>*Nat Methods* (2023) | Zhao, C. et al.<br>*Nat Methods* (2023) | Zhao, C. et al.<br>*Opt Express* (2023) | Zong, W. et al.<br>*Cell* (2022) | Madruga, B.A. et al.<br>*Nat Commun* (2025) | This work |
| --- | --- | --- | --- | --- | --- | --- | --- |
| **Lens architecture** |  |  |  |  |  |  |  |
| **Excitation**<br>**Collection** |  |  |  |  |  |  |  |
| **Modality (2p/3p)** | 3p | 3p | 3p | 2p | 2p | 2p | 2p |
| **Collection NA** | 0.9 | 0.9 | 0.65 | 0.45 | 0.45-0.5 | 0.6 | 0.95 |
| **Excitation NA** | 0.48-0.54 | 0.48 / 0.58 | 0.55 | 0.25 | 0.45-0.5 | 0.36 | 0.26 |
| **Lateral resolution** | 0.8 $\mu\text{m}$ | 1.18 / 1.11 $\mu\text{m}$ | 0.97-1.21 $\mu\text{m}$ | 1.47 $\mu\text{m}$ | 1.15-1.24 $\mu\text{m}$ | 0.98 $\mu\text{m}$ | 0.97-0.99 $\mu\text{m}$ |
| **Axial resolution**<br>**(depth of field)** | 5.3 $\mu\text{m}$ | 13.8 / 10.1 $\mu\text{m}$ | 7.21-8.51 $\mu\text{m}$ | 24.64 $\mu\text{m}$ | 12.8-17.8 $\mu\text{m}$ | 10.18 $\mu\text{m}$ | 80.37-90.23 $\mu\text{m}$ |
| **FOV** | 160 × 160 $\mu\text{m}^2$ | 300 × 300 /<br>200 × 200 $\mu\text{m}^2$ * | 400 × 400 $\mu\text{m}^2$ | 1000 × 788 $\mu\text{m}^2$ | 420 × 420 /<br>500 × 500 $\mu\text{m}^2$ ** | 445 × 380 $\mu\text{m}^2$ | 423 × 439 $\mu\text{m}^2$ |
| **Frame rate** | 27.78 Hz<br>@ 120 × 120 pixels | 10.6 Hz<br>@ 273 × 280 pixels | 15.93 / 8.35 Hz<br>@ 128 × 128 /<br>200 × 200 pixels | 9 Hz<br>@ 600 × 512 pixels | 40 / 15 Hz<br>@ 256 × 256 pixels | 8.62 / 7.92 Hz<br>@ 512 × 354 /<br>512 × 350 pixels | 9 Hz<br>@ 512 × 512 pixels |
| **Pixel rate** | 0.4 megapixels/s | 0.8 megapixels/s | 0.3 megapixels/s | 2.8 megapixels/s | 2.6 megapixels/s | 1.6 megapixels/s | 2.4 megapixels/s |

Note: Major developments after 2020 with a known lens structure were listed here.
\* Two configurations: extended z-range with 0.48 NA, 300 × 300 $\mu\text{m}^2$ FOV; high-resolution with 0.58 NA, 200 × 200 $\mu\text{m}^2$ FOV.
\*\* Two configurations: MINI2P-F with 420 × 420 $\mu\text{m}^2$ FOV, 40 Hz frame rate; MINI2P-L with 500 × 500 $\mu\text{m}^2$ FOV, 15 Hz frame rate.

**Supplementary Table 2:** Major parameters used for miniBB2p imaging.

| | Indicator | Paradigm | Microscope | FOV ($\mu\text{m}^2$) | Frame rate | Imaging duration | 3D data projection* | Laser power (mW)** | Imaging depth ($\mu\text{m}$) |
| --- | --- | --- | --- | --- | --- | --- | --- | --- | --- |
| **Fig. 2f (Left)** | Thy1-GFP | Brain slice | miniBB2p | 400 × 400 | 9 Hz | 11.1 s (100 frames) | mean | 80 | 40 ± 40 |
| **Fig. 2f (Right)** | Thy1-GFP | Brain slice | Tabletop 2P | 400 × 400 | 1.9 Hz | 14.7 s (28 images) | sum | 25 | 0~81 |
| **Fig. 2g (Left)** | Thy1-GFP | Brain slice | miniBB2p | 60 × 60 | 9 Hz | 11.1 s (100 frames) | mean | 80 | 40 ± 40 |
| **Fig. 2g (Right)** | Thy1-GFP | Brain slice | Tabletop 2P | 60 × 60 | 0.16 Hz | 176 s (28 images) | sum | 25 | 0~81 |
| **Fig. 2h, Video S1** | GCaMP7f | Free-moving | miniBB2p | 423 × 439 | 9 Hz | 6 min 10 s | s.d. | 40 | 185 ± 40 |
| **Fig. 2k** | GCaMP7f | Head-fixed | Tabletop 2P | 423 × 439 | 0.8 Hz | 35 s (28 images) | sum | 12 | 145~226 |
| **Fig. 2l** | GCaMP7f | Head-fixed | Tabletop 2P | 423 × 439 | 7.7 Hz | 6 min (2 min/layer) | mean | 9 | 145, 185, and 226 |
| **Fig. 4a Video S5** | GCaMP7f | Free-moving | miniBB2p | 420 × 420 | 9 Hz | 30 min | s.d. | 40 | 160 ± 40 |
| **Fig. 5b Video S6** | GCaMP7f | Free-moving | miniBB2p | 420 × 420 | 9 Hz | 8 min | s.d. | 40 | 150 ± 40 |
| **Fig. S5a** | Thy1-GFP | Head-fixed | miniBB2p | 420 × 420 | 9 Hz | 11.1 s (100 frames) | mean | 80 | 80 ± 40 |
| **Fig. S5b** | Thy1-GFP | Head-fixed | Tabletop 2P | 420 × 420 | 1 Hz | 28 s (28 images) | sum | 25 | 40~121 |
| **Fig. S5c** | Thy1-GFP | Head-fixed | miniBB2p | 120 × 120 | 9 Hz | 11.1 s (100 frames) | mean | 80 | 80 ± 40 |
| **Fig. S5d** | Thy1-GFP | Head-fixed | Tabletop 2P | 120 × 120 | 1 Hz | 28 s (28 images) | sum | 25 | 40~121 |
| **Fig. S6a Video S2** | GCaMP8s | Head-fixed | miniBB2p | 420 × 420 | 9 Hz | 27 s | mean | 80 | 80 ± 40 |
| **Fig. S6b Video S3** | GCaMP8s | Head-fixed | miniBB2p | 60 × 60 | 9 Hz | 3 min | mean | 100 | 100 ± 40 |
| **Fig. S7a Video S4** | GCaMP8s | Free-moving | miniBB2p | 120 × 120 | 9 Hz | 30 min | mean | 80 | 110 |
| **Fig. S8a** | GCaMP7f | Head-fixed | miniBB2p | 420 × 420 | 9 Hz | 30 min | s.d. | 40 | 170 ± 40 |
| **Fig. S9a (Bottom)** | GCaMP7f | Head-fixed | Tabletop 2P | 420 × 420 | 1 Hz | 31 s (31 images) | sum | 10 | 130~220 |
| **Fig. S11b** | GCaMP8s | Free-moving | miniBB2p | 420 × 420 | 9 Hz | ~4 min | mean | 60 | 145 ± 40 |

\* Various z-projection methods were used for the presentation of the 3D data
\*\* Laser power is measured after the objective lens

**Supplementary Table 3:**
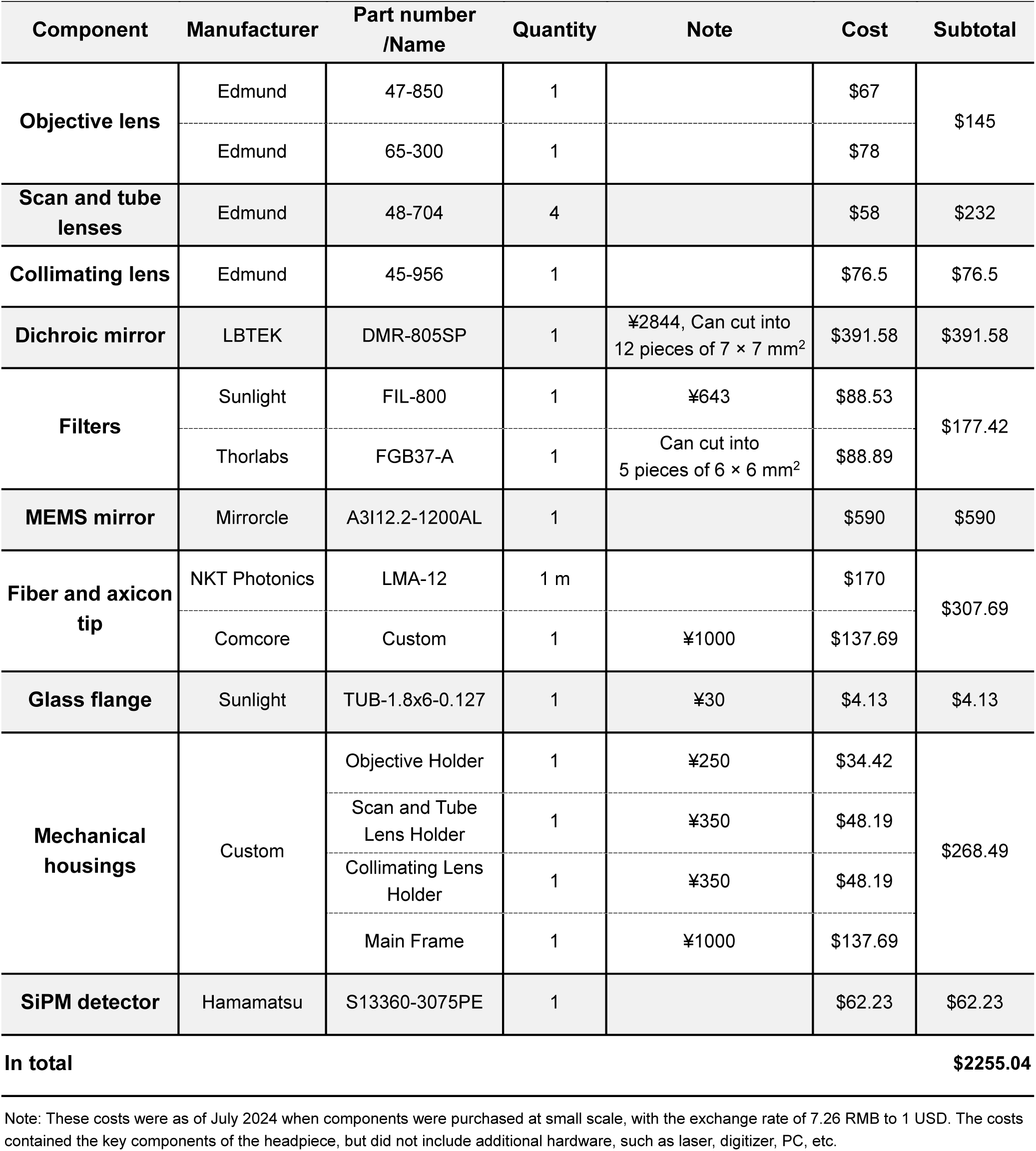
Cost of the miniBB2p headpiece.

## SUPPLEMENTARY VIDEO INFORAMTION

**Supplementary Video 1, Imaging the neural activities in ACC in a freely moving mouse.** Images were captured at 9 frames per second, 423 × 439 µm^2^ FOV, and 40 mW post-objective power. Data were corrected for motion artifacts, binned every 3 frames, and saved at 25 binned frames per second (top left). Behavior video of the mouse in the chamber during imaging (top right). Calcium traces of example neurons (bottom).

**Supplementary Video 2: Imaging the activities of dendritic trees in ACC in a head-fixed awake mouse.** Images were captured at 9 frames per second, 420 × 420 µm^2^ FOV, and 80 mW post-objective power. Data were corrected for motion artifacts and saved at 25 frame per second.

**Supplementary Video 3: Imaging the activities of dendrites and spines in ACC in a head-fixed awake mouse.** Images were captured at 9 frames per second, 60 × 60 µm^2^ FOV, and 100 mW post-objective power. Data were corrected for motion artifacts, binned every 3 frames, and saved at 25 binned frames per second.

**Supplementary Video 4: Imaging the activities of dendrites and spines in ACC in a freely moving mouse over a 30-minute period.** Images were captured at 9 frames per second, 120 × 120 µm^2^ FOV, and 80 mW post-objective power. Data were corrected for motion artifacts (smoothness parameter set to [2, 2] in *Flow-Registration*), binned every 3 frames, displayed at accelerated speeds (×75 for first 15 minutes and ×8.3 for final 15 minutes), and saved at 25 binned frames per second. Behavior video of the mouse in the chamber during imaging (top right). Calcium traces of example dendrite and spines (bottom).

**Supplementary Video 5: Imaging the neural activities in M2 in a freely moving mouse over a 30-minute period.** Images were captured at 9 frames per second, 420 × 420 µm^2^ FOV, and 40 mW post-objective power. Data were corrected for motion artifacts, binned every 9 frames, and saved at 25 binned frames per second (top left). Behavior video of the mouse in the chamber during imaging (top right). Calcium traces of example neurons (bottom left) and movement trajectory of the mouse (bottom right).

**Supplementary Video 6: Imaging the response of ACC neurons to pain stimulus in a freely moving mouse.** Images were captured at 9 frames per second, 420 × 420 µm^2^ FOV, and 40 mW post-objective power. Data were corrected for motion artifacts, binned every 3 frames, and saved at 25 binned frames per second (left). Behavior video of the mouse under non-noxious and noxious stimulus (right).

